# Repeated horizontal acquisition of lagriamide-producing symbionts in Lagriinae beetles

**DOI:** 10.1101/2024.01.23.576914

**Authors:** Siddharth Uppal, Samantha C. Waterworth, Alina Nick, Heiko Vogel, Laura V. Flórez, Martin Kaltenpoth, Jason C. Kwan

## Abstract

Microbial symbionts associate with multicellular organisms on a continuum from facultative associations to mutual codependency. In some of the oldest intracellular symbioses there is exclusive vertical symbiont transmission, and co-diversification of symbiotic partners over millions of years. Such symbionts often undergo genome reduction due to low effective population sizes, frequent population bottlenecks, and reduced purifying selection. Here, we describe multiple independent acquisition events of closely related defensive symbionts followed by genome erosion in a group of Lagriinae beetles. Previous work in *Lagria villosa* revealed the dominant genome-eroded symbiont of the genus *Burkholderia* produces the antifungal compound lagriamide and protects the beetle’s eggs and larvae from antagonistic fungi. Here, we use metagenomics to assemble 11 additional genomes of lagriamide-producing symbionts from seven different host species within Lagriinae from five countries, to unravel the evolutionary history of this symbiotic relationship. In each host species, we detected one dominant genome-eroded *Burkholderia* symbiont encoding the lagriamide biosynthetic gene cluster (BGC). Surprisingly, however, we did not find evidence for host-symbiont co-diversification, or for a monophyly of the lagriamide-producing symbionts. Instead, our analyses support at least four independent acquisition events of lagriamide-encoding symbionts and subsequent genome erosion in each of these lineages. By contrast, a clade of plant-associated relatives retained large genomes but secondarily lost the lagriamide BGC. In conclusion, our results reveal a dynamic evolutionary history with multiple independent symbiont acquisitions characterized by high degree of specificity. They highlight the importance of the specialized metabolite lagriamide for the establishment and maintenance of this defensive symbiosis.

## Introduction

Eukaryotes have been associated with prokaryotic microbes since the initial endosymbiotic events that led to the acquisition of mitochondria and chloroplasts [1]. These organelles represent the presumed endpoint of ancient symbioses with α-proteobacteria and cyanobacteria, respectively, that over time led to a progressive shrinkage of the symbionts’ genomes and eventual transfer of genes from symbionts to host [1]. Although organelle acquisition appears to be a rare event [2], other more recent symbioses appear to be on a similar evolutionary trajectory of profound genome reduction and absolute dependence on host cells. For example, the acquisition of the intracellular symbiont *Buchnera aphidicola* in the common ancestor of aphids allowed them to diversify as sap-feeding insects since the symbiont synthesises several essential amino acids not found in plant sap, as evidenced by a rapid basal radiation of aphid species [3] and strict co-evolution of aphids and *Buchnera* [4]. *B. aphidicola* has been vertically transmitted for at least 200 million years [4] and has a profoundly reduced chromosome, about 11% of the size of *Escherichia coli* [5]. Through comparison of various symbionts, a model of genome reduction has emerged whereby host-restriction initially weakens purifying selection on formerly essential genes, through both host-provided metabolites and symbiont population structure, with low effective population sizes and isolation within individual hosts [6]. When symbionts are vertically transmitted, population bottlenecks occur during every transmission event, causing the fixation of deleterious mutations within the population [6]. These factors combine to first cause an increase in pseudogenes in the genome [6] and then deletion of those pseudogenes due to a known deletion-bias within bacteria [7]. The most reduced genomes lose even central functions such as DNA repair pathways [6], which leads to an increased rate of evolution and further gene loss, as well as increased AT-bias in many cases [8, 9]. In the cases of symbionts living inside host cells, it is likely that this process is exacerbated due to a lack of opportunity or ability to horizontally acquire functional genes.

However, genome reduction is also known to occur without genetic isolation. For instance, free-living bacteria living in nutrient-poor environments such as *Prochlorococcus* spp. are thought to have reduced genomes as a consequence of selection pressure to streamline their metabolism [10], potentially explained through the Black Queen hypothesis [11], which posits that selection drives pathways to be lost when they are produced by another species in the ecosystem as “public goods”. There are also genome-reduced symbionts which seemingly are not genetically isolated. *Burkholderia* symbionts that reside extracellularly in leaf nodules in plants are mainly transmitted vertically because the symbiosis is mutually co-dependent [12], although horizontal transfer may have occurred rarely between plants, the soil microbiota, and insects [13]. This suggests a lack of genetic isolation, and indeed there is evidence of repeated horizontal transfers of biosynthetic genes for defensive molecules among leaf nodule symbionts of *Rubiacaeae* plants [13]. Such systems may provide an opportunity to study the evolutionary pressures that lead to the process of genome reduction, and the mechanisms of symbiosis that underlie it.

The dichotomy of vertical versus horizontal transfer of symbionts may be one determinant of genome reduction. A relatively clear-cut example is the two symbionts of the tunicate *Lissoclinum patella*, the extracellular cyanobacterium *Prochloron didemni* [14] and the intracellular “*Candidatus* Endolissoclinum faulkneri” [15]. The former is capable of horizontal transmission, which is reflected in its almost clonal genome amongst very divergent hosts and a lack of genome reduction [14], while the latter is vertically transmitted, as evidenced by its co-divergence with its hosts across cryptic speciations, and profound genome reduction [15, 16]. However, the mode of transmission also exists on a continuum from strict vertical to strict horizontal, with mixtures of vertical and horizontal transmission in between [17]. For instance, the Tsetse fly symbiont *Sodalis glossinidius* shows some signs of genome-reduction such as rampant pseudogenes, but remains culturable in the lab, meaning that horizontal transmission cannot be excluded [18]. Likewise, symbionts long thought to be exclusively vertically transmitted, such as the bryozoan symbiont “*Ca.* Endobugula sertula”, which is packaged with the hosts’ larvae, show no signs of genome reduction [19], indicating that there is no compelling reason why it should not be able to transmit horizontally between hosts. Indeed, “*Ca*. E. sertula” has been found in genetically divergent but proximal bryozoan individuals, suggesting horizontal transmission [20].

The *Lagria* and *Ecnolagria* beetles belong to subfamily Lagriinae within the family Tenebrionidae (order Coleoptera). The *Lagria villosa* beetle, a known soybean pest [21], is a source of lagriamide, an antifungal polyketide [21]. The compound is produced via a *trans*-AT polyketide synthase (PKS)-non-ribosomal peptide synthetase (NRPS) hybrid biosynthetic gene cluster (BGC), termed *lga,* which due to a nucleotide signature (kmer frequency) distinct from the chromosome is predicted to have been horizontally acquired [21]. The *lga* BGC is encoded by a *Burkholderia* symbiont (*Burkholderia* sp. LvStB) that is present in glandular structures associated with the ovipositor of female beetles and secreted on its eggs as they are laid [21]. The symbiont has been shown to have a defensive role against fungi in the egg [21] and larval stages [22]. Previously, we showed that the genome of *Burkholderia* sp. LvStB is reduced and has lost several essential genes including some genes involved in the DNA repair pathways and primary metabolism [23]. The genome has a low coding density, and a high number of pseudogenes and transposases, indicative of a reduced genome [23]. These characteristics are consistent with host restriction and vertical transmission of LvStB. However, there is evidence that *Burkholderia* symbionts from *L. villosa* can be transferred to plant tissues and survive for several days and that bacteria can be acquired by the beetle from the plant and soil environment [24].

As some Lagriinae beetles harbor symbionts in special structures that likely evolved between 55 and 82 million years ago based on fossil evidence, positioned to deposit symbionts on the eggs [25, 26], we hypothesized that lagriamide-producing *Burkholderia* symbionts might have co-evolved with their hosts in a manner similar to other vertically-transmitted insect symbionts. However, the possibility for transmission of the symbionts to and from plants, and the accessibility of the symbionts’ habitat on the surface of eggs and within adult females suggested that horizontal symbiont acquisition may be possible. As the beetles harbor complex microbiomes with multiple related *Burkholderia* strains as well as other bacteria [22, 24], both genome-reduced and not, an alternative hypothesis is that the lagriamide BGC has been repeatedly horizontally transferred among environmental strains and symbionts. Moreover, partnerships in defensive symbionts are usually more dynamic as compared to intracellular nutritional symbionts [27]. It is also possible that the lagriamide-producing strain is restricted to *L. villosa*, and that different Lagriinae species have symbionts with different BGCs, as this would allow the association to react much more flexibly to changes in antagonist communities. To clarify this evolutionary picture, we analyzed the metagenomes of 12 beetle samples, spanning seven species belonging to the genera *Lagria* and *Ecnolagria* across five different countries (four continents) (**Table 1**). We recovered the metagenome-assembled genomes (MAGs) of several different *Burkholderia* bacteria and confirmed the presence of the lagriamide BGC in each specimen. We also report a complete genome of the genome-reduced, lagriamide-producing *Burkholderia* sp. LvStB symbiont, obtained through long-read Nanopore sequencing. We compared the phylogeny of the recovered *Burkholderia* MAGs, the lagriamide BGCs, and the host beetles to determine whether co-cladogenesis occurred in this system, and to further explore the evolutionary relationships in the symbiosis. The results indicate that the lagriamide BGC was likely only acquired once in the common ancestor of beetle-associated *Burkholderia* symbionts, and subsequently lost in the majority of the descendent free-living strains. Remarkably, as all the lagriamide-bearing symbionts are genome-reduced but do not form a monophyletic clade and do not correspond to host phylogeny, they likely represent multiple acquisition events, followed by independent genome-reduction processes. The common factor of lagriamide production might be one of the reasons for selection by and dependency on hosts. This would suggest that a single group of natural products caused several independent symbioses to be established over evolutionary time.

**Table 1.**
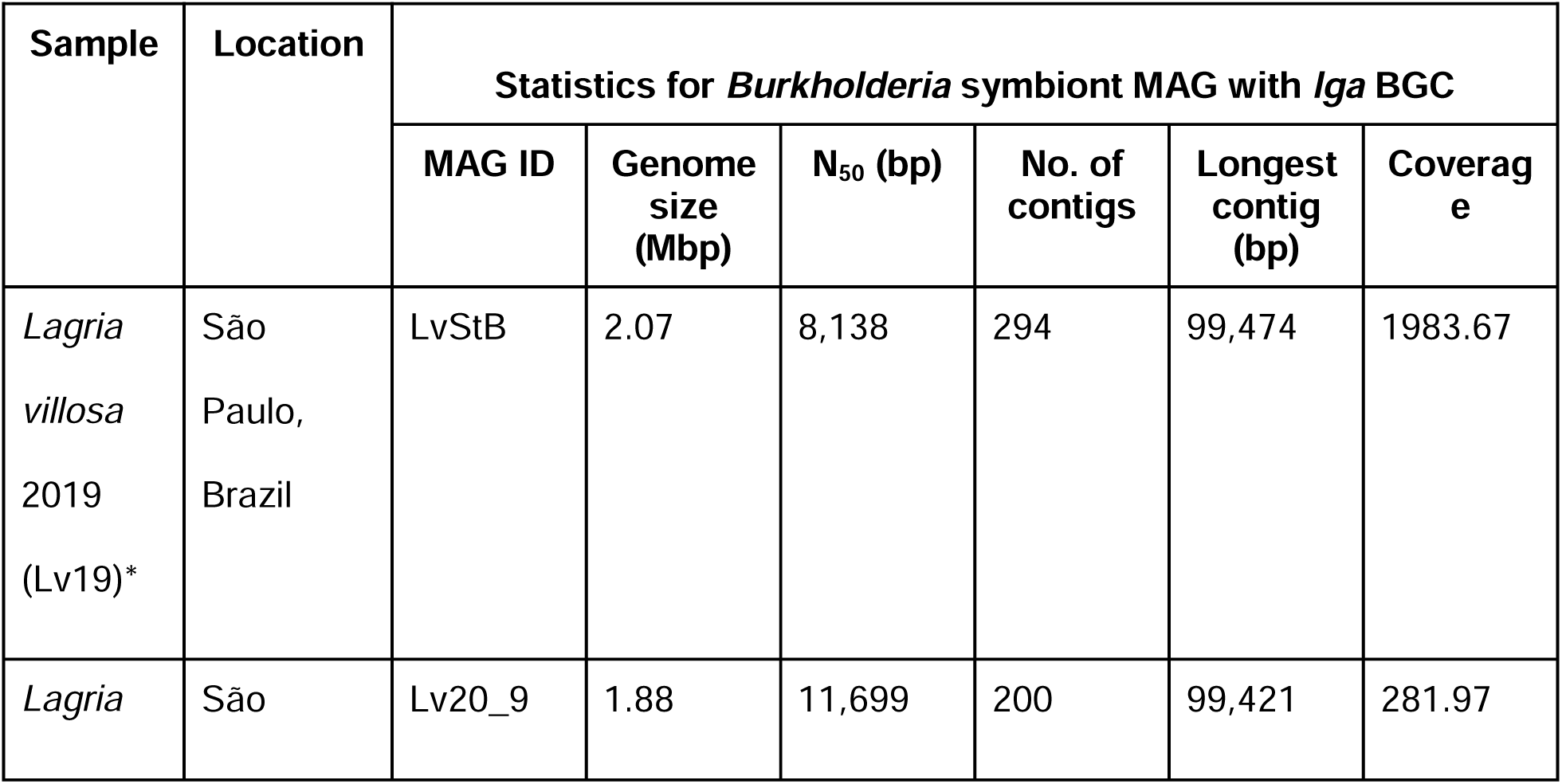
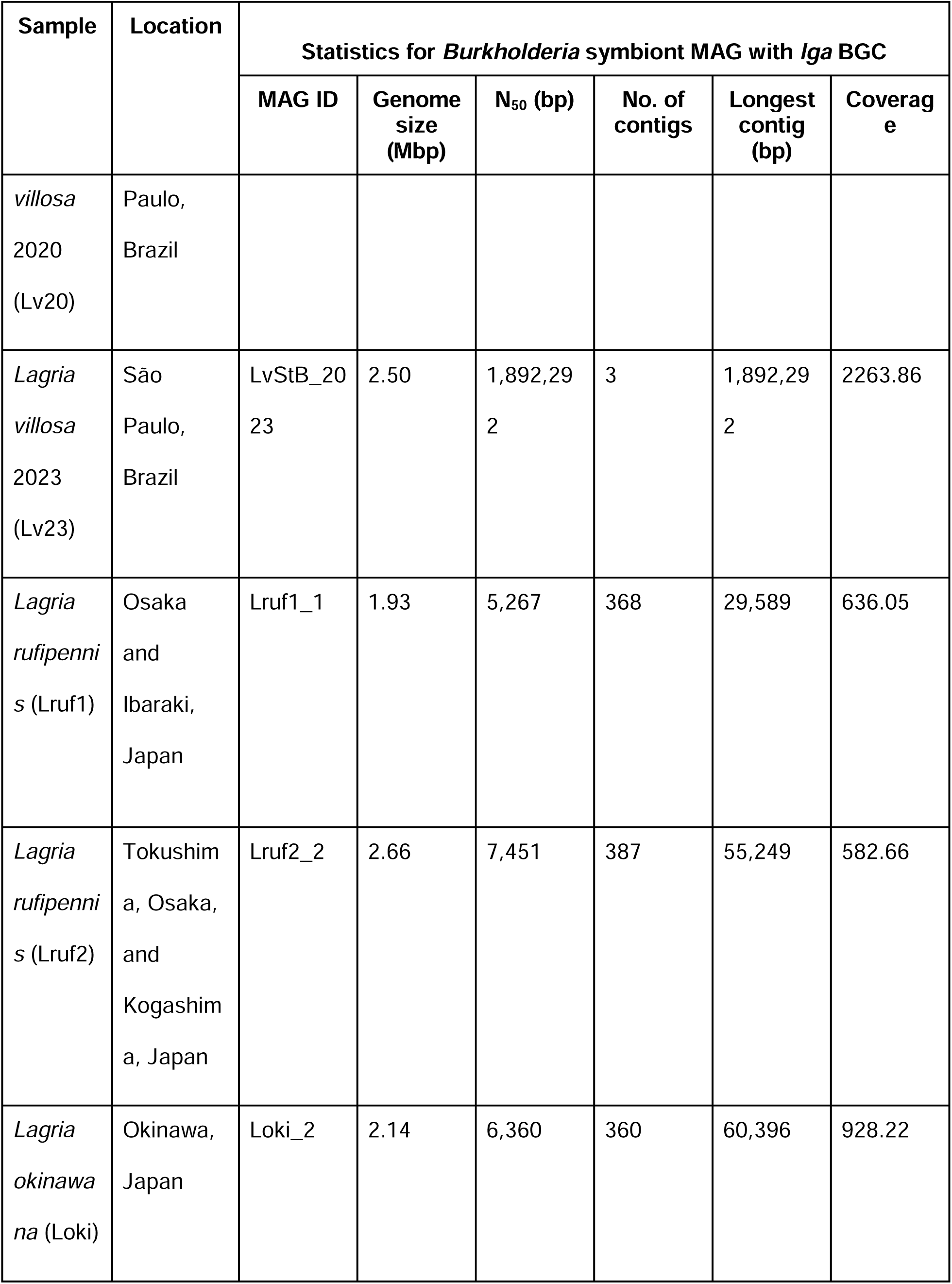
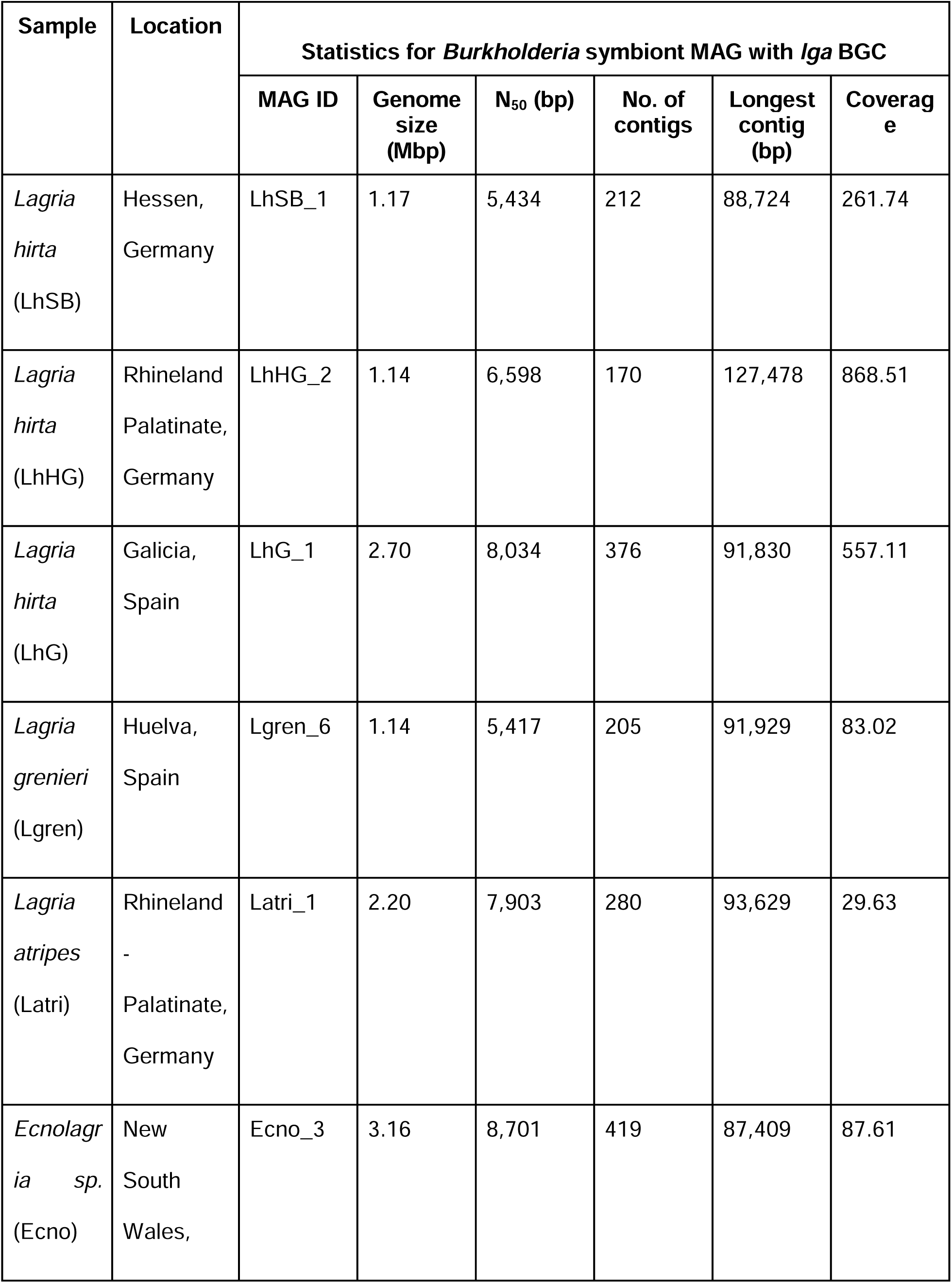
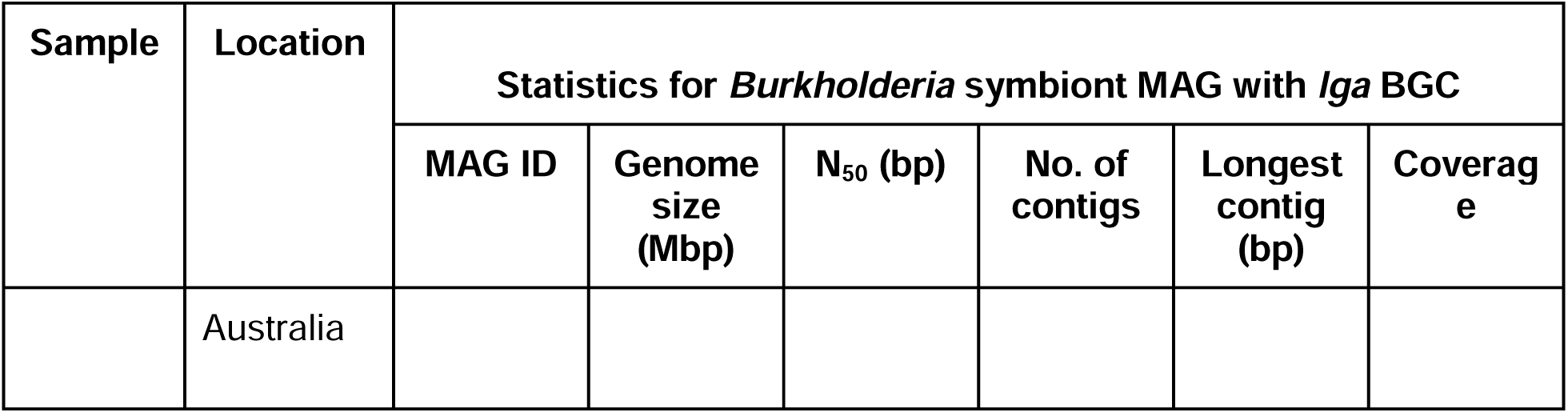
Metadata for different beetles collected for this study. \**Lagria villosa* samples were collected at three different time points, with one sample reported in a previous study (referred to as Lv19 in this work) [21, 23].

## Results and Discussion

### Beetle phylogeny

We sequenced and assembled the metagenomes of the collected *Lagria* and *Ecnolagria* beetle populations (**Table 1**), and beetle mitogenomes (see **Table S1** for mitogenome statistics) were extracted and annotated to infer host beetle phylogeny (**Fig. 1**). In line with previous studies, mitogenomes belonging to the tenebrionid subfamilies Lagriinae, Blaptinae, Pimeliinae, Stenochiinae, and Alleculinae were found to be monophyletic whereas Diaperinae and Tenebrioninae were found to be para- or polyphyletic [28–31]. Maximum likelihood analysis using RAxML [32] (**Fig. SI 1**) and Bayesian analysis using MrBayes [33] (**Fig. 1**) gave similar results.

**Figure 1.**
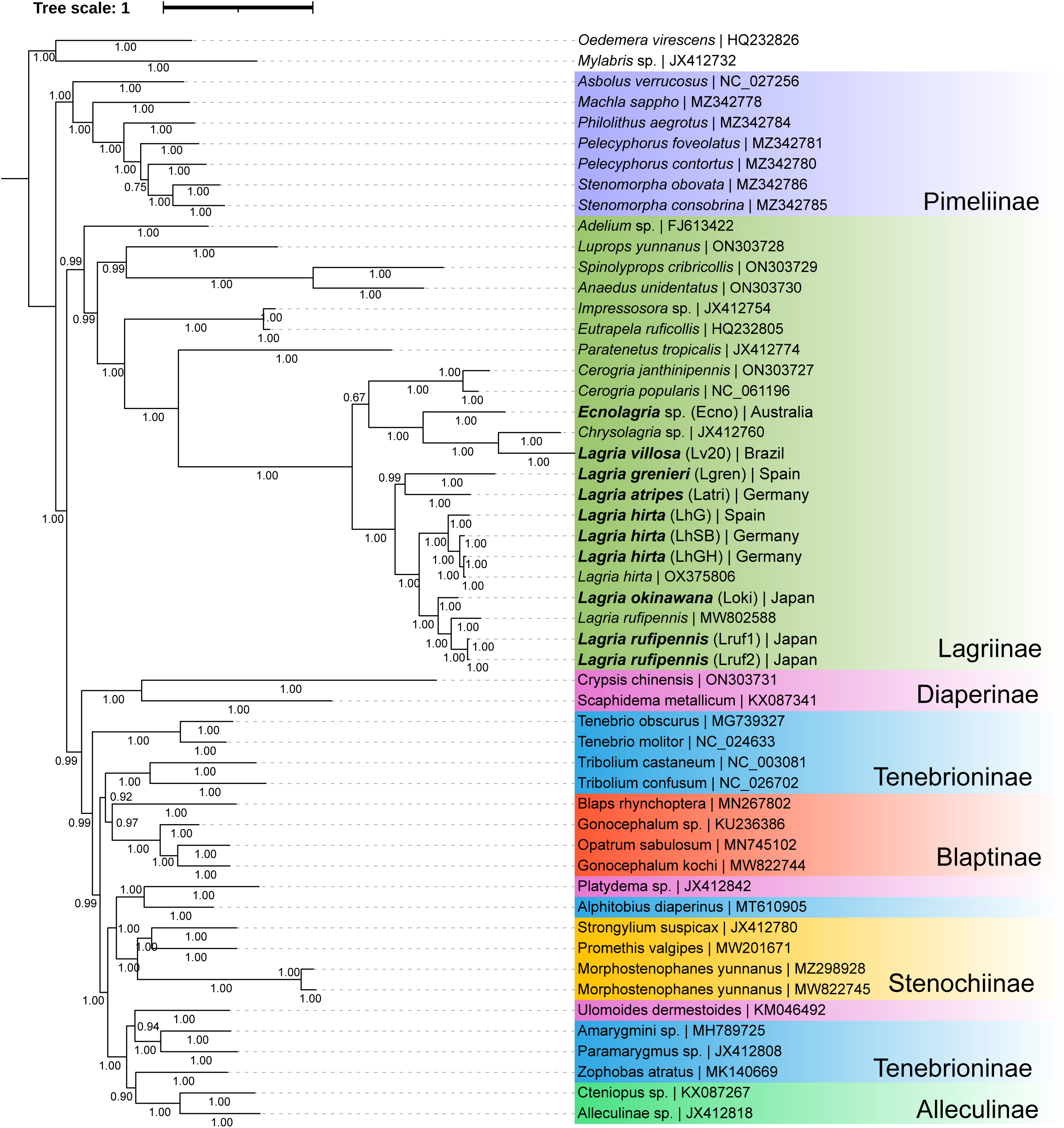
Beetle mitogenome phylogenetic tree using 13 mitochondrial protein coding genes constructed using MrBayes [33]. Branch values represent posterior probabilities. Mitogenomes recovered in this study are highlighted with bold lettering.

All collected *Lagria* beetle mitogenomes clustered into four distinct subclades: All *L. hirta* beetle mitogenomes were clustered in a single clade, alongside a closely related clade of *Lagria* species (*L. rufipennis* and *L. okinawana*) from Japan. The *L. atripes* and *L. grenieri* beetles formed another clade more distantly related to the *L. hirta* and Japanese *Lagria* species. Finally, the *L. villosa* and *Ecnolagria* sp. beetles formed a fourth clade along with *Chrysolagria* sp. (JX412760), distinct from the other *Lagria* beetles. A small distinction was noted here, wherein the Bayesian phylogeny (**Fig. 1**) suggested that the *Cerogria* beetles belonged to the clade with *L. villosa* and *Ecnolagria* species, whereas the Maximum-likelihood phylogeny showed the

*Cerogria* to be in a clade with all other *Lagria* beetles (**Fig. SI 1**). However, in both cases the branch support values are too low to make any definite conclusions. Publicly available sequences of *L. hirta* (OX375806) clustered with collected *L. hirta* samples from Rhineland Palatinate, Germany (LhHG), and *L. rufipennis* (MW802588) clustered with the two *L. rufipennis* (Lruf1 and Lruf2) samples.

### Recovery of Lagriamide BGCs

A complete, or mostly complete, *lga* BGC was found, using antiSMASH v7 [34, 35], in eleven of the twelve samples, with the exception of *L. rufipennis 1* where only small fragments of the *lga* BGC could be recovered. The BGC recovered from *L. rufipennis 2* was found over two contigs and could not be manually joined following inspection of the assembly graph. The missing data for this region spans from approximately halfway through the *lgaB* gene to approximately halfway through the *lgaC* gene (**Fig. 2A**).

**Figure 2.**
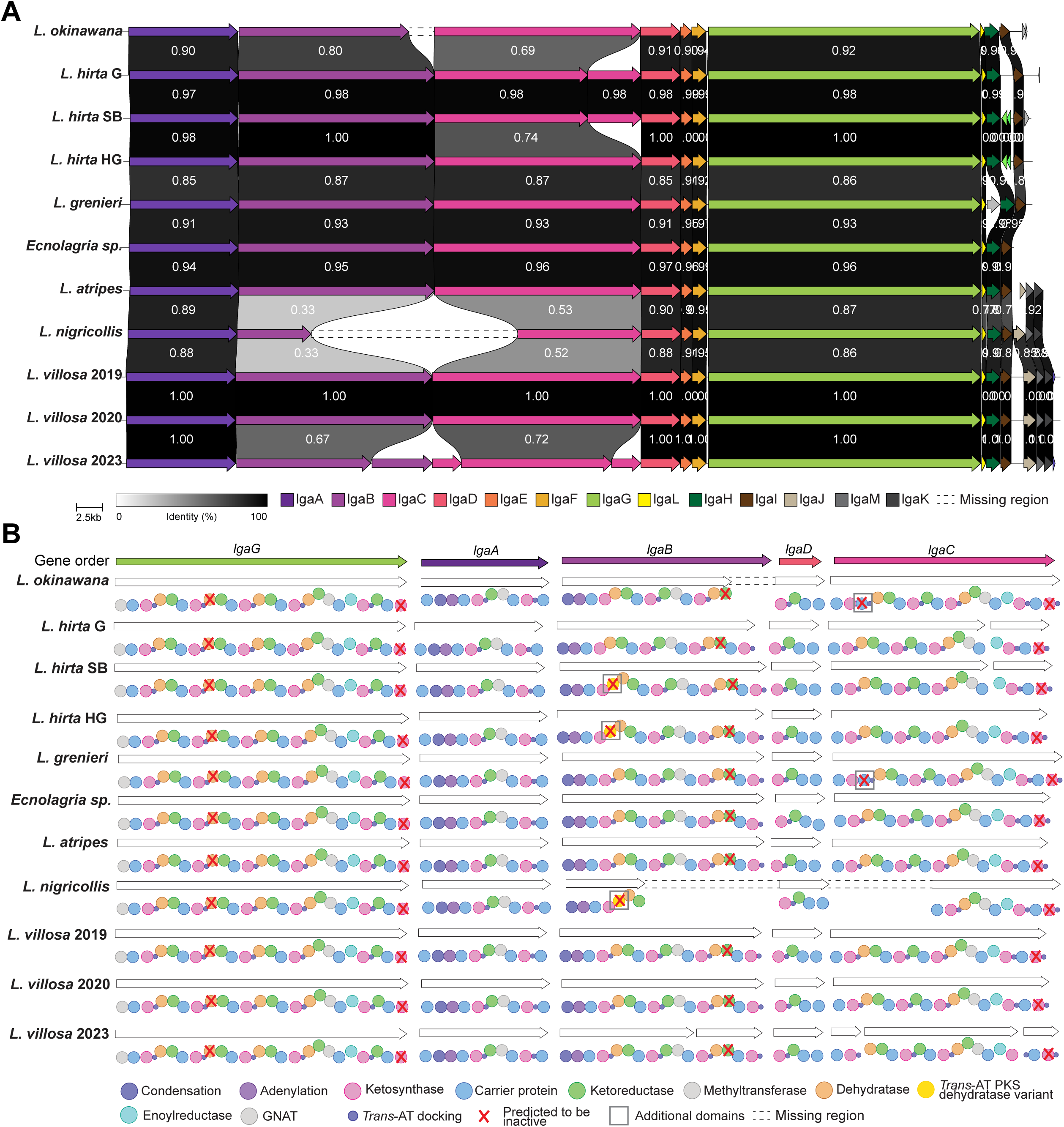
Analysis of representative *lga* BGCs extracted from eleven Lagriinae beetle metagenomes. A) Comparison of representative *lga* BGC gene organization. Individual genes in the *lga* BGCs are represented by arrows oriented in the predicted direction of transcription and colored according to identity. Pairwise amino acid similarity between BGCs is indicated in the shaded areas between genes. A scale bar is provided for gene size. Dashed lines indicate fragments missing from the respective assemblies. B) Comparison of predicted enzyme domain organization in the representative *lga* BGCs, where genes are ordered according to biosynthetic order. Boxes around the domains indicate differences between the BGCs.

Analysis of representative BGCs revealed two differences in gene organization of the *lga* BGC across the different Lagriinae beetle species (**Fig. 2A**). The first difference observed regarded the *lgaC* gene. The *lgaC* gene from the BGCs recovered from the *L. hirta* G and *L. hirta* SB samples appeared to be split into two, denoted *lgaC1* and *lgaC2* for clarity. Alignment of the *lgaC* gene from the three *L. hirta* samples revealed that there was a perfect alignment of the nucleotide sequences save for a 37-bp deletion in the BGCs from *L. hirta* G and *L. hirta* SB, which introduced a frameshift (**Fig. SI 2A**). This frameshift consequently introduced a premature stop codon which split the *lgaC* gene into two ORFs in *L. hirta* G and *L. hirta* SB (**Fig. SI 2B**).

The second difference we observed was in the *lga* BGC from the LvStB genome extracted from the 2023 *L. villosa* metagenome (Lv23). One split in *lgaB* and two splits in *lgaC* were seen.

However, the assembly of the *L. villosa* 2023 metagenome was based solely on long-read data, which is error-prone [36], and the splits may not be a true reflection of the BGC in this sample. Normally Sanger sequencing would be the solution to validate these questionable regions but unfortunately, there was no remaining DNA after the long read sequencing runs for this particular sample. For this reason, we left the BGC with the splits but were cautious not to over-interpret the apparent breaks in the genes in this BGC.

We then considered the domain organisation within the *lga* BGC genes (**Fig. 2B**). The domain organisation is largely congruent across the *lga* BGCs recovered from the metagenomes. We did note, however, an additional annotated “DHt” domain in *lgaB*, which is defined as “Dehydratase domain variant more commonly found in *trans*-AT PKS clusters”, in the *lga* BGCs from all *L. hirta* samples and the *L. rufipennis 2* sample. Similarly, we detected an additional carrier protein domain (phosphopantetheine acyl carrier protein group) near the N-terminus of the *lgaC* protein in the BGCs from the *L. grenieri* and *L. okinawana* samples. In all cases, close inspection of the primary sequence of these additional dehydratase and carrier domains revealed mutations in the sequences that would likely render the encoded domain non-functional (see Supplementary Material for full details).

Finally, as with the originally described *lga* BGC recovered from the *L. villosa* 2019 sample [21], we found mutations in the catalytic or conserved motifs of *lgaG* DH2, *lgaG* KS6, *lgaB* KR3 and *lgaC* KS5 domains, that we believe may render these domains inactive (see Supplementary Material for full details). As a result, the domain architecture of all representative *lga* BGCs from all samples appear functionally identical.

Together, the conservation of the *lga* BGC in at least seven different species of Lagriinae beetles, across four geographically distant countries, implies that the production of lagriamide is an important factor for the host beetle and that the *lga* BGC is under strong selective pressure. The presence of additional domains in the *lga* BGC in several samples, even though they are likely inactive, is intriguing as it suggests that these domains may have previously been present in all *lga* BGCs but may have decayed over time and were lost. The reason as to why these domains were selected against would be speculative at best and inspection of all lagriamide-like compounds produced in the different beetle populations would need to be assessed to truly infer differences that the domain architecture may have on the resulting chemistry. Conserved production of other bioactive compounds has been observed, such as pederin, across Staphylinidae beetle species (*Paederus* and *Paederidus* genera) [37], which are host to a *Pseudomonas* symbiont that produces pederin [38].

The two systems have several parallels: both pederin and lagriamide are produced by a *trans*-AT PKS NRPS hybrid BGC in *Pseudomonas* bacterium [37], where the compound is concentrated in the female oviposition organs, coated onto the eggs and serves to protect juveniles [39]. Further, both pederin and lagriamide are the sole insect-associated compounds in suites of compounds otherwise associated with marine invertebrates. Groups of pederin analogs, such as the onnamides, mycalamides, psymberins, and theopederins have been isolated from a variety of marine sponges [40–44] and ascidians [45], while bistramide, the most structurally similar compound to lagriamide, was isolated from an ascidian [46]. The question remains, however, as to what the fundamental link is between these terrestrial and marine systems.

### Complete genome of the *lga*-carrying LvStB symbiont

Long-read sequencing of the *L. villosa* 2023 (Lv23) metagenome allowed us to assemble a complete genome of a *lga*-carrying *Burkholderia* strain (referred to as LvStB_2023 from hereon). LvStB_2023 was found to have a 2.5 Mbp long genome with a GC percentage of 58.63%. It has 2 circular chromosomes - chromosome 1 is 1.89 Mbp, chromosome 2 is 0.55 Mpb in size, and there is a plasmid 59.77 kbp long. The genome is estimated to be 97.1% complete (98.8% with “specific” mode) and 0.02% contaminated as per CheckM2 [47] and thus is a high-quality MAG according to the MIMAG standards [48]. Assembly graph analysis of LvStB_2023 verified that we have the complete sequence of two circular chromosomes and a plasmid. However, the CheckM2 estimate did not reflect a fully complete genome, at 98.8%, and we believe that this small discrepancy in predicted completeness may be a result of ongoing genome reduction [49].

LvStB_2023 has a coding density of 78% and 59.1% with and without pseudogenes respectively. A large percentage (43.87%) of the ORFs in LvStB_2023 were identified as pseudogenes (1613 out of 3676), the highest of any *lga*-carrying *Burkholderia* symbiont. However, this estimate may be artificially high as pseudogenes are identified purely based on their length relative to their closest BLASTP match and these counts are derived from an assembly generated from only long-read data which can be prone to errors [50–52], particularly homopolymeric runs. However, coding density and frequency of pseudogenes is not very different from LvStB MAGs assembled from short-read data (see **Table S2** for complete genome characteristics of recovered MAGs). Having multiple chromosomes is a common phenomenon in *Burkholderia* [53, 54]. Generally in multi-chromosome bacteria, the majority of the genes for essential functions are located on one larger or primary chromosome, while the smaller or secondary chromosome has much fewer essential genes and mostly carries genes for niche specific functions [55]. In the case of LvStB_2023, chromosome 1 appears to be the primary chromosome as it is much larger in size, and has 77 out of 84 core genes (including multiple copies) (**Fig 3A**). Functional analysis revealed chromosome 1 to have the highest number of genes for all essential COG categories (**Fig 3B**), including categories L (replication, recombination and repair), J (Translation, ribosomal structure and biogenesis), M (Cell wall/membrane/envelope biogenesis) and H (Coenzyme transport and metabolism).

**Figure 3.**
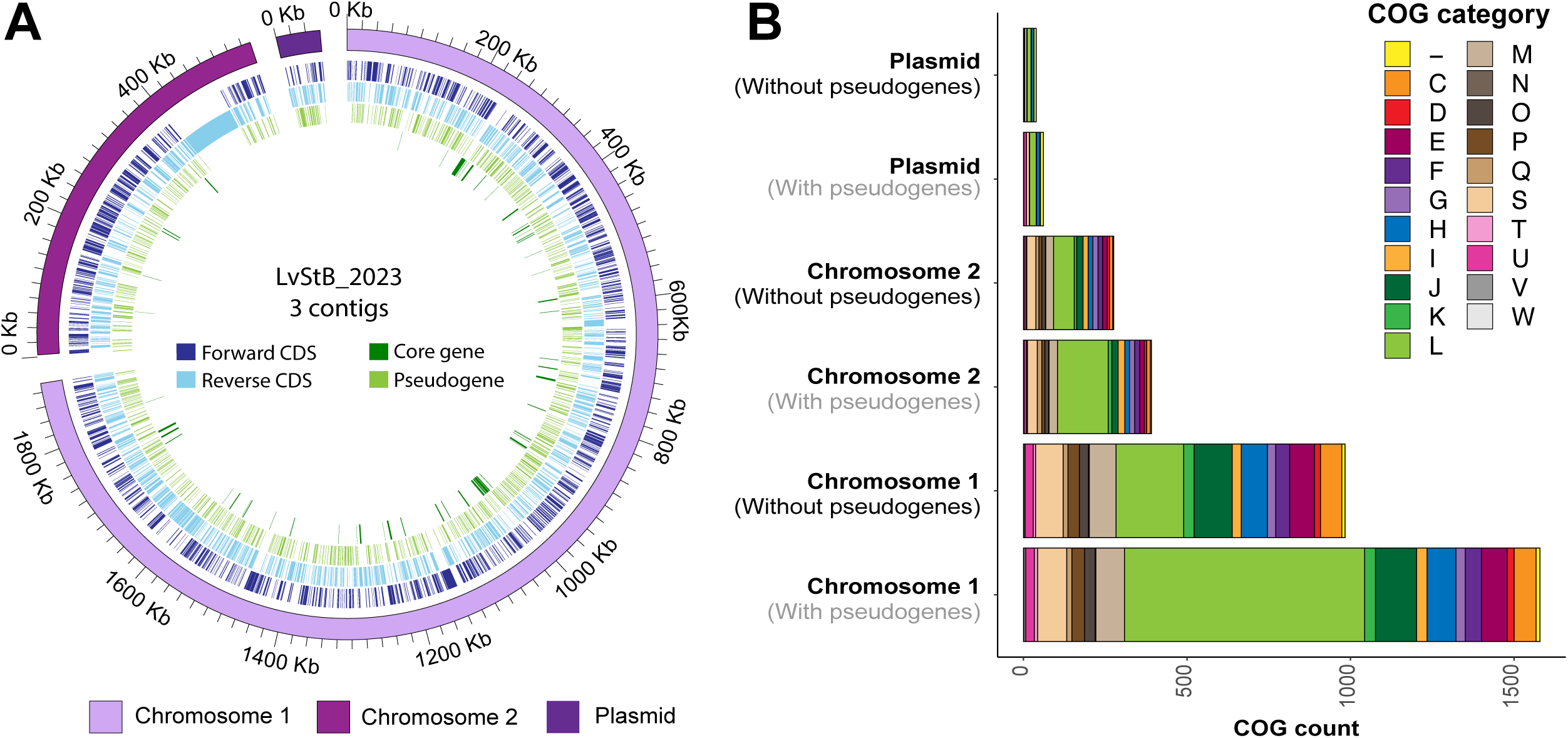
A) Circular representation of LvStB_2023 genome from the *L. villosa* 2023 sample. Individual chromosomes are indicated by shades of purple. Coding sequences (CDS) which are core genes or pseudogenes, as indicated by shades of green, while the rest are indicated in shades of blue. B) Raw count of COG categories present on different contigs of the LvStB_2023 genome (with and without pseudogenes) from the *L. villosa* 2023 sample.

The *lga* BGC is on chromosome 2 (0.55 Mbp long) and can be distinguished by the continuous block of coding sequences on the reverse strand in **Fig. 3A**. A detailed view of the *lga*-carrying chromosome can be seen in **Fig. SI 3,** where the GC content of the *lga* BGC is significantly higher than the rest of the genome, providing further evidence for its horizontal acquisition. Chromosome 1, chromosome 2, and the plasmid have 44.75%, 37.59%, and 42.34% of their coding capacity taken up by pseudogenes, respectively. The similar abundance of pseudogenes in each of the contigs indicates that the whole genome is undergoing reduction simultaneously. The chromosome with *lga* (chromosome 2) has the smallest percentage of pseudogenes, which may be a reflection of the required conservation of the *lga* BGC in combination with the presence of large genes in *lga*.

### Diversity of beetle-associated *Burkholderia* symbionts

#### Recovery and analysis of metagenome-assembled genomes

Following assembly, the 12 beetle metagenomes were binned, and the resultant bins were manually refined. A total of 77 MAGs were recovered from all samples, of which 24 MAGs were of high quality, 30 of medium quality, and 23 of low quality (**Table S2**) in accordance with published MIMAG standards [48]. Only medium and high-quality MAGs were used for downstream analysis, with the exception of one low-quality bin carrying the *lga* BGC (LhHG_2). Confirmation that the *lga* BGC had been assigned to the correct MAG has been covered in detail in the Supplementary Material. Genome erosion, such as that already observed for the *lga*-carrying symbiont *Burkholderia* sp. LvStB [23], can skew the completeness metric. To determine if a lower quality MAG was incomplete or genome-reduced, we also considered several other metrics, including core gene presence, number of pseudogenes [23], and coding density (**Table S2**), and concluded that this particular MAG (LhHG_2) was likely both reduced and incomplete.

For each beetle population, a single MAG belonging to the genus *Burkholderia* with a single copy of the *lga* BGC was identified. Previous studies on the lagriamide-carrying symbiont strain *B. gladioli* LvStB [21, 23, 56], showed that this strain was significantly more abundant than all other bacteria associated with *L. villosa*, and had a reduced genome. Consistent with this, all newly recovered MAGs that included *lga* BGCs were the most abundant MAGs in each sample, had reduced genomes with an abundance of pseudogenes and transposases, and had lower coding densities relative to other *B. gladioli* genomes (**Table S2**). In standing with previous studies of *Lagria* beetles, where both reduced and non-reduced *B. gladioli* genomes were recovered, additional *B. gladioli* MAGs (Latri_2, LhHG_3, and LhSB_5) were recovered that did not carry the lagriamide BGC and showed no evidence of genome erosion. We also recovered three small *B. gladioli* MAGs (Lgren_7, Lv19_6_18, Lv20_2) and one small *Burkholderia* MAG (Lv19_6_14), as well as MAGs classified as *B. lata* (Lv19_4_0) and *B. arboris* (Lv20_1).

Average nucleotide identity (ANI) analysis of *B. gladioli* MAGs carrying the *lga* BGC showed that MAGs from different beetle species and/or different locations were likely different bacterial species due to shared ANI values less than 95% [57]. However, previous studies have suggested that ANI alone is not a sufficient metric for species delineation and that the aligned fraction (AF) must also be taken into account [57–60]. Following recent cutoffs adopted for species delineation [58], we opted to use AF ≥ 60% along with ANI ≥ 95% as a cutoff for species assignment. Subsequently, we found that the *Burkholderia* MAGs carrying the lagriamide BGC appeared to be split into at least five novel species (**Table S3**).

#### Phylogenetic analysis of recovered metagenome-assembled genomes

In order to elucidate the evolutionary history of the association between Lagriinae beetles and *Burkholderia* symbionts, we reconstructed phylogenies of the *Burkholderia* symbionts and free-living relatives based on shared single-copy genes. A priori, we hypothesized that the *lga*-encoding, genome-eroded symbionts would form a monophyletic clade Showing co-diversification with the hosts, given that such patterns have been previously described across many ancient and co-evolved symbioses.

The phylogeny of the beetle-associated *Burkholderia* symbionts, relative to other *Burkholderia* species, was inferred using 126 single-copy hierarchical orthogroups (HOGs) (non-pseudogenes) present in more than 90% of the genomes (RAxML phylogeny is represented in **Fig. 4** and **Fig. SI 4** while **Fig. SI 5** represents the Bayesian approach). *Burkholderia* symbionts without the *lga* BGC were broadly present across the phylogeny containing *B. gladioli, B. lata,* and *B. arboris* strains. By contrast, and consistent with our expectation, symbionts of different host species carrying the *lga* BGC were closely related. Surprisingly, however, these genome-eroded, *lga*-encoding symbionts did not form a monophyletic clade. Because the tree indicates that the common ancestor of the genome-reduced *lga*-encoding symbionts also gave rise to a lineage of non genome-reduced descendents, this result indicates a non-reduced free living common ancestor and subsequent multiple independent acquisition events by Lagriinae beetles. To test for the robustness of our phylogenetic analysis, we repeated the analysis using single-copy HOGs present in 95%, 80%, 70%, and 60% (**Fig. SI 6**) of the genomes, as well as after removing any putative horizontally transferred genes (**Fig. SI 7 and 8**). Other than minor discrepancies in the terminal nodes, we obtained highly similar phylogenetic trees, supporting the lack of monophyly of the *lga* BGC carrying *Burkholderia* symbionts. Thus, all our analyses support a phylogeny that contains a clade of mostly free-living (plus some beetle-associated symbionts with non-eroded genomes) *Burkholderia* that groups within the *lga* BGC-containing Lagriinae symbionts. Concerning the evolutionary history of the symbiosis, this leaves us with two alternative scenarios: (i) an ancestral association of the whole clade of bacteria with beetles and a certain degree of genome erosion on the deep branches, and a subsequent reversal to a free-living stage of the clade containing many plant-associated *B. gladioli* strains; or (ii) at least four independent transitions from a free-living (or plant-associated) to a symbiotic lifestyle, each of which followed by genome erosion.

**Figure 4.**
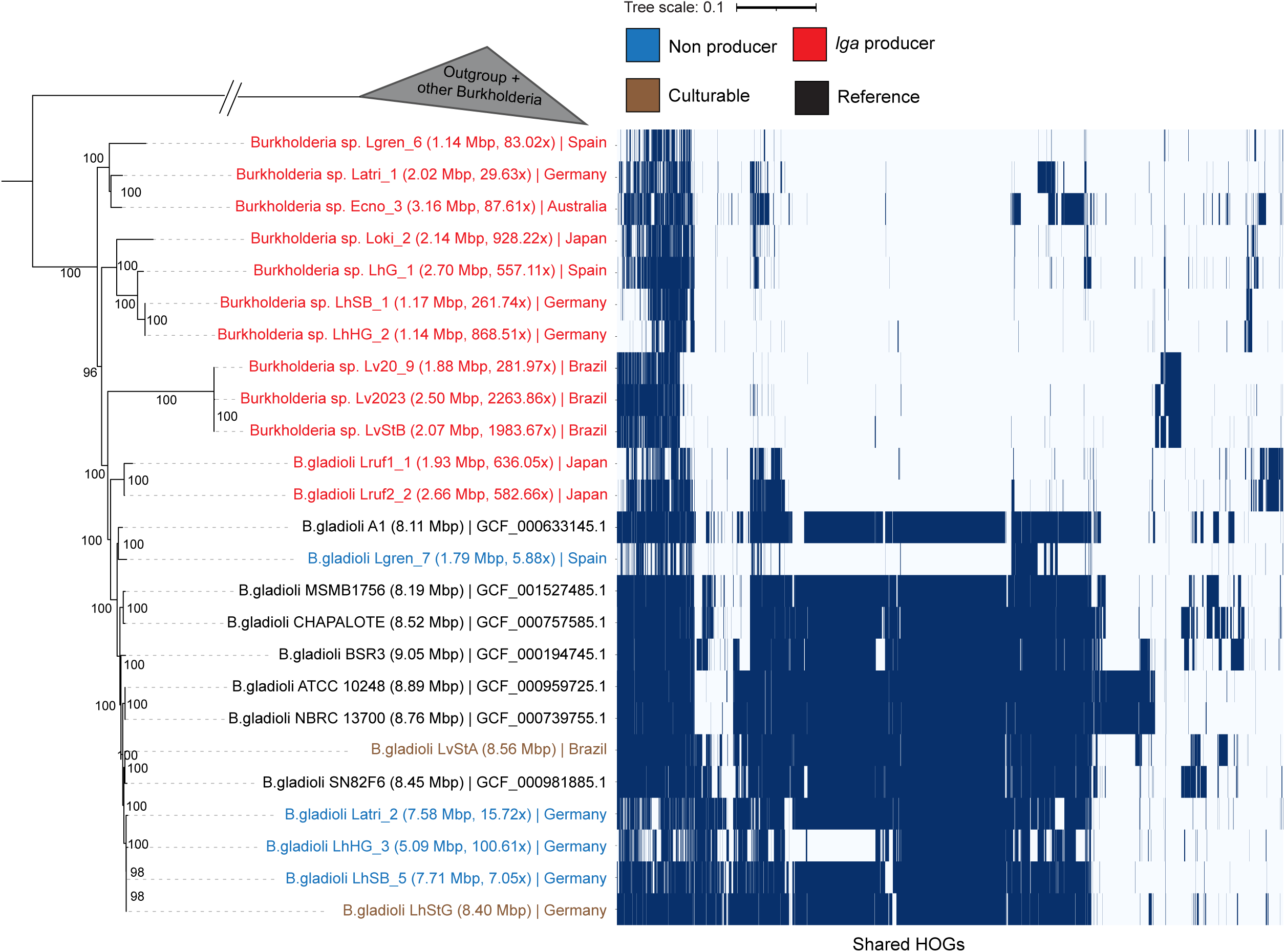
RAxML phylogenetic tree (left) and shared hierarchical orthogroups (HOGs) (non-pseudogenes) between different *Burkholderia* genomes (matrix on the right). Each blue line indicates a shared HOG. HOGs have been hierarchically clustered on the x-axis to improve visualization. Values on nodes indicate bootstrap values. Genome size and coverage is represented in brackets next to MAG ID. Outgroups include - *Paraburkholderia acidiphila* (GCF_009789655.1), *Cupriavidus necator* (GCF_000219215.1), *Herbaspirillum seropedicae* (GCF_001040945.1). Tree has been truncated for visualisation purposes, full tree can be found in **Fig. SI 4** (RAxML phylogeny) and **Fig. SI 5** (bayesian phylogeny).

To unravel which of these scenarios is more likely, we analyzed shared HOGs between different *Burkholderia* sp., after removing any pseudogenes from MAGs. We observed substantially higher conservation of orthogroups between the potentially free-living *Burkholderia* sp. than among the *lga*-containing symbionts, with the free-living strains sharing a large core genome (**Fig. 4**). If the shared ancestor of all *lga*-encoding symbionts and the free-living strains would have been tightly associated with beetles and experienced some degree of genome erosion (scenario i), this observation would postulate a substantial increase in the genome size of the bacteria after the reversal to the free-living/plant-associated lifestyle and before the clade split into the different taxa. While theoretically possible, this scenario seems highly unlikely, because acquisition of a large number of genes would have to have happened quickly and early in order for extant strains in this clade to have such a degree of gene overlap. Instead, it appears much more plausible that there were at least four independent transitions to a symbiotic lifestyle, each of which was followed by genome erosion. This is consistent with the observation that the genomes of the eroded strains retain distinct sets of genes, many of which represent subsets of the free-living strains’ core genomes (**Fig. SI 9**), since gene loss from independant host-restriction events would be expected to be largely stochastic. This distinct set of genes can however, also be due to symbiont replacement events followed by genome reduction.

Furthermore, the lack of synteny observed in the genes flanking the *lga* BGC (**Fig. SI 10**) is indicative of either genomic rearrangement or independent acquisition of the *lga* BGC. Both of these further support the independent acquisitions of symbionts followed by genome erosion. It is important to note that this conclusion is based on the current data and may change as we obtain more samples and long-read metagenomes that allow for synteny analyses across the entire genome.

Consistent with the scenario of multiple independent transitions to a symbiotic lifestyle, the phylogeny of the *lga* BGC-carrying *Burkholderia* symbionts was found to be largely incongruent with the beetle phylogeny (**Fig 5A**), except for the symbionts grouping together for individuals of the same host species, i.e. *L. hirta* and *L. rufipennis*, respectively. The incongruence between host and symbiont phylogenies suggests both multiple symbiont acquisition and possibly host switching events that lead to symbiont replacements. Symbiont replacement has often been reported in nutritional symbionts as a way for the hosts to replace a genetically degraded symbiont with a more complete and effective one and to acquire new adaptations for expanding into different niches [61]. *Burkholderia* symbionts related to *B. gladioli* in *Lagria* beetles have been reported to evolve from plant-associated bacteria [26] capable of transfer from beetles to plants with subsequent survival [24]. It is possible that the horizontal acquisition might occur in the egg and larval stages, where the symbionts are localized on the surface (eggs) or in cuticular invaginations (larvae and pupae) that remain connected to the external surface via a small duct [62]. As the closely related *Burkholderia* strain LvStA can be acquired horizontally from the environment [24], and there is evidence of free-living bacteria carrying lagriamide-like BGCs [63], we propose that there are *lga*-carrying *Burkholderia* strains persevering in the environment (e.g. in plants or soil) [24] that can be horizontally acquired by the beetle host.

**Figure 5.**
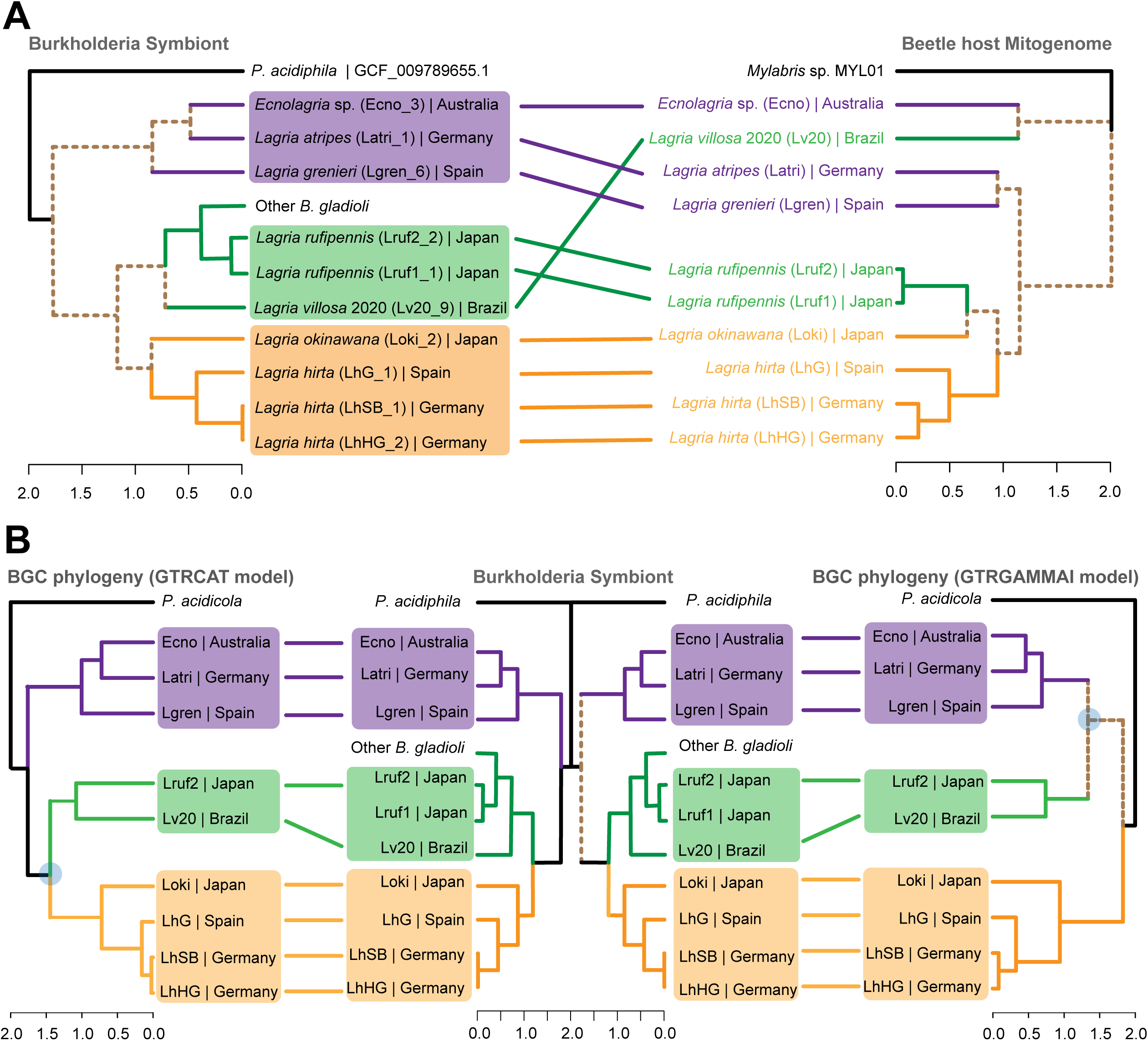
Congruence between phylogenies of beetle host, *Burkholdria* symbionts and *lga* BGCs in all samples. A) Tanglegram between *lga*-carrying symbionts and beetle host phylogeny. B) Tanglegram between *lga*-carrying symbionts (centre) and the *lga* BGC, as inferred via two models GTRCAT (left) and GTRGAMMAI (right). Clades within the beetle host mitogenome phylogenetic tree are highlighted in shades of grey. In all panels, the three conserved clades are highlighted in purple, green and orange. Dashed brown lines represent nodes that are unique between the respective phylogenies, except when the node is absent from the other tree. Blue circles indicate clade nodes that are poorly supported.

As we previously observed that *lga* has distinct nucleotide composition to the Lv19 genome [21], suggestive of a recent horizontal transfer, we sought to determine if it has been independently transferred to the corresponding symbiont in different beetle hosts. Phylogenetic analysis of the representative *lga* BGCs from all samples resulted in two possible topologies using GTRCAT-V and GTRGAMMAI models (**Fig. 5B**). Both topologies included conserved clades. However, the relative positions of the three clades are poorly supported (**Fig. 5B**, highlighted in blue), resulting in the two alternative topologies. A Bayesian tree was also constructed (**Fig. SI 11**) which is congruent with the GTRGAMMAI tree topology. The inconsistent topology likely stems from limited resolution of the phylogeny affecting deep nodes in the trees. The GTRCAT-V topology is perfectly congruent with the symbiont phylogeny based on genome-wide marker genes, while the GTRGAMMAI topology shows one discrepancy at one of the deep nodes. Thus, these analyses do not provide evidence for additional horizontal transfer events of the *lga* cluster, so it is likely that there was a single acquisition of *lga* in the common ancestor of the symbiont and *B. gladioli* clade, with subsequent loss in the free-living group. It appears that lagriamide production was highly selected for in symbiotic settings and hence retained, whereas it was lost in the most of the larger genomes (assumed to be free-living) where it was not selected for. These findings indicate that the *lga* BGC is important in the symbiosis, either for symbiont establishment (e.g. competition with other symbionts) and/or because lagriamide is an effective host-defensive molecule. Furthemore, the fact that highly similar *Burkholderia* species with *lga* were identified across different Lagriinae beetles indicates that symbiont acquisition is highly selective.

Interestingly, lagriamide seems to be highly conserved, despite the dynamics of the system, where multiple species of bacteria associate with each beetle host, and several *lga*-producing *Burkholderia* have apparently been independently acquired. A dynamic association in defensive symbioses has been previously hypothesized, to allow for rapid adaptation to a changing community of antagonists, or to individual co-evolving pathogens [27]. We expected to see changes in the defensive chemistry used in a symbiotic context, akin to the rapid evolution of immune genes in animals [64–66]. However, despite the dynamic nature of many defensive symbioses, with symbiont replacements on ecological or evolutionary timescales, several examples of defensive symbioses highlight that the same defensive compounds can be used over long evolutionary timescales. In case of beewolf wasps, *Streptomyces* symbionts have been found to produce piericidin and streptochlorin for an estimated 68 million years [67, 68]. Both compounds are found in different beewolf species and across different geographic locations. Similarly, as discussed above, pederin is produced across different species of *Paederus* and *Paederidus* beetles by *Pseudomonas* symbionts [37, 38]. Similarly, we now describe the production of lagriamide by a *Burkholderia* symbiont across several species of *Lagria* and *Ecnolagria* beetles. Thus, even though these defensive symbioses are dynamic in the acquisition and replacement of microbial partners, the chemistry seems to be conserved. This suggests a limited diversity of chemical compounds that can be used for defense against eukaryotic antagonists (predators or fungi) in a symbiotic context, which is supported by the convergence on similar compounds in terrestrial and aquatic symbioses. It is possible that this might be due to the harmful side effects of the bioactive molecule on the eukaryotic host, analogous to the cytotoxic side-effects of antifungal pharmaceuticals on humans, resulting in only limited diversity of such compounds.

To gain insights on the possible origin of the *lga* BGC, we performed an analysis of pentanucleotide (5-mer) frequencies of the beetle-associated, *lga*-carrying symbionts and their associated BGCs, along with the genomes of recently identified soil-borne *Paraburkholderia* species that carry the lagriamide B (*lgb)* BGC, which is highly similar to the *lga* BGC [63]. Visualisation of 5-mer frequencies of the BGCs and the genomes revealed three clusters of BGCs: The BGCs from the two soil-borne *Paraburkholderia* strains, the BGCs from the Brazilian *L. villosa*-derived LvStB strains, and then a third cluster of all other *lga* BGCs (**Fig. SI 12**). A similar pattern was observed for the nucleotide composition of the respective genomes wherein LvStB and Lv20_9 form an isolated cluster, the two soil-borne *Paraburkholderia* form a second, distant cluster, and all other *lga*-carrying *Burkholderia* strains and cultured *Lagria*-associated genomes (LvStA and LhStG) form a third cluster. None of the BGCs share similar 5-mer composition with their respective genomes, providing additional evidence for the horizontal acquisition of the *lga* BGC.

We noted during the analysis of the COG annotated genes in LvStB_2023 that there appeared to be a particularly high number of genes in the L category (replication, recombination and repair) that were likely pseudogenes (**Fig 3B**). We assessed the percentage change of COG annotated genes in all *lga*-carrying *Burkholderia* and found that this pseudogenization of genes involved in DNA replication, recombination and repair was particularly high in all the Brazilian *L. villosa*-derived LvStB strains, as well as the MAGs LhSB_1, LhG_1, Loki_2 and Lgren_6 (**Fig. SI 13**). Two of the three LvStB strains also exhibited high pseudogenization of the genes associated with cell motility (Category N). While COG annotation of genes does not provide a robust picture, as not all genes are successfully annotated, the increased pseudogenization of genes involved in DNA replication and repair may explain the divergence of the LvStB strains observed in both the phylogenetic analysis and the related 5-mer analysis. In particular, LvStB MAGs possessed highly truncated and psuedogenized *polA* genes, coding for DNA polymerase I used in many DNA-repair pathways and chromosome replication [69], whereas other *lga*-containing MAGs, except LhHG_2, had intact *polA* genes (**Fig. SI 14**). The loss of *polA* in the *L. villosa* symbionts explains their accelerated sequence evolution in the genome as a whole and also in the *lga* BGC compared to other *lga*-possessing symbionts (**Fig. SI 12**). The absence of *polA* in LhHG_2 could be due to its poor quality, as it is only 46% complete and has only 47.6% percentage of core genes.

Previous studies have highlighted how symbionts can be conserved across host-speciation events and millions of years, leading to genome reduction in the symbiont [6, 16]. A disadvantage of such an exclusive relationship is that the symbiont inevitably suffers from increasing genome erosion that can result in reduced efficiency in providing benefits to the host [70]. Consequently, many long-term obligate symbioses have experienced symbiont replacement events that can provide an escape route for the insect host after its symbiont entered the irreversible phase of degenerative genome reduction [71]. Such replacement is a common phenomenon in Hemipteran symbionts [61]. In the present study, however, we are suggesting that the repeated replacement of symbionts may have happened with very closely related strains that carry the same biosynthetic gene cluster and hence likely provide the same functional benefit to the host. One reason we are suggesting multiple acquisitions and displacements may have happened is that all the *lga*-containing symbionts appear to be at different stages of genome reduction, with very different genome sizes and gene complement, perhaps indicating that they have been symbionts for different amounts of time. That in combination with the apparent importance of *lga* specifically, the incongruence of symbiont and host phylogeny, and the fact that none of the symbionts is profoundly genome-reduced, suggests that although Lagriinae likely hosted *lga-*containing symbionts since the evolution of special symbiont storage structures, the current symbionts are not direct descendents. The replaced symbionts were likely genome-reduced to an extent that they were outcompeted by incoming *lga-*bearing strains from the environment. The *lga* BGC-containing *Burkholderia* strains were consistently the most abundant symbionts in the metagenomes across seven different Lagriinae species, indicating that the *lga* BGC or an as yet unknown genomic feature shared among the symbiont strains provides a key selective advantage in the beetles’ symbiotic organs. Possibly, lagriamide is uniquely suited to defend the symbionts’ niche against competitors and/or protect its host from antagonists. However, as lagriamide shows lower antifungal activity than some secondary metabolites of related *Burkholderia* strains [21, 26, 72], another intriguing possibility is that it only provides a moderate degree of defence but at the same time exhibits less harmful side effects on the host than other antifungal compounds. Further elucidating the relevance of lagriamide in establishing the symbiotic association with beetles will not only provide valuable insights into the ecological and evolutionary dynamics of defensive symbioses, but may also unravel the mechanisms ensuring specificity in symbiotic alliances.

## Methods

For additional details see Supplementary Material.

### Insect collection

Specimens were collected between 2009 and 2019 in Spain, Germany, Brazil, Japan and Australia in the locations listed in **Table S4**. Female adults were dissected either directly after chilling for ca. 15 min at –20°C or preserved in 70% ethanol or acetone until dissected. The accessory glands were removed and preserved in 70% ethanol at –80°C until further processing. For species in which we suspected the presence of symbiont-harbouring compartments within the ovipositor in addition to the glands, the ovipositor was also dissected and preserved along with the accessory glands.

### DNA isolation and metagenomic sequencing

Genomic DNA from the preserved organs was extracted per individual after removing the fixative and homogenizing the tissue in liquid nitrogen. The MasterPure^TM^ complete DNA and RNA isolation Kit (Epicentre Technologies) was used as indicated by the manufacturer, including an additional incubation step at 37°C for 30 min with 4 µL lysozyme (100 mg mL^-1^) before protein precipitation. The nucleic acids were re-suspended in Low TE buffer (1:10 dilution of TE) and pooled by species. Metagenomic sequencing was carried out in two batches. The first batch included the samples corresponding to *L. atripes*, *L. grenieri*, *L. hirta* G, *L. hirta* SB, *L. hirta* HG, and *L. villosa* 2020. This first batch was sent for DNA library preparation using a Nextera XT DNA Library Prep Kit (Illumina) and metagenomic sequencing on a NovaSeq 6000 platform, using a paired-end approach (2 x 150 bp) to a depth of 30 M reads (9 Gbp) by CeGaT GmbH (Tübingen, Germany). Samples from the second batch including *L. rufipennis 1 and 2*, *L. okinawana*, and *Ecnolagria sp.*, were sequenced using Illumina NextSeq 2000 (paired end 2 x 150bp) to a depth of 28 to 44 million reads at the Max Planck Genome Centre (Cologne, Germany). The data from sample Lv19 corresponds to that described in Florez *et al*. [21]. Taxonomic assignment of individual specimens of *L. rufipennis* was first done morphologically according to Jung and Kim [73] very similar to the sympatrically occurring females of *Lagria nigricollis*, and the specimens used for sample Lruf2 were originally identified as *L. nigricollis* (see also [26]). Due to the uncertainty associated with morphological identification, we therefore additionally barcoded the specimens after their metagenomes had been sequenced (see Supplementary information for detailed methods) and compared their cytochrome oxidase I sequences to those of male specimens of *L. rufipennis* and *L. nigricollis* that can be more easily distinguished based on their morphology. All 19 *L. rufipennis* and 10 *L. nigricollis* COI sequences that we obtained turned out to be very similar and formed a sister group to the *L. rufipennis* sequence available in NCBI (MW802588). However, the *L. nigricollis* sequences formed a distinct subclade, with the exception of the sample that had been used for metagenomics (Lruf2), which grouped within *L. rufipennis*. Hence, we reassigned Lruf2 to *L. rufipennis*, resulting in two replicate metagenomes for this species. Unfortunately, the *L. nigricollis* samples were males (which do not contain symbionts), preventing us from sequencing a metagenome of this species.

For the *L. hirta* HG population, genomic DNA was extracted from a pool of 6 egg clutches (20-30 eggs per clutch) using the Qiagen Genomic-tip 20/G Kit (Qiagen) following the instructions from the manufacturer. The sample was split in half for DNA extraction and the resulting genomic DNA was again combined. This sample and an aliquot of the *L. villosa* 2020 genomic DNA sample underwent End-DNA repair and library preparation using the NEBNext Ultra II DNA Library Prep Kit (New England Biolabs, Ipswich, USA) and the Ligation Sequencing Kit V14 (SQK-LSK114; Oxford Nanopore Technologies, Oxford, UK) followed by a clean-up step with AMPure XP beads (Beckman Coulter). These were sequenced on a MinION platform (Oxford Nanopore Technologies) as described below for long-read sequencing of the *L. villosa* 2023 sample.

### Metagenomic sequence assembly and binning

Sample *L. villosa* 2019 represented data previously assembled and analyzed [21, 23]. Sequence data generated from *L. atripes*, *L. grenieri*, *L. hirta* G, *L. hirta* SB, *L. rufipennis 1*, *L. rufipennis 2*, *L. okinawana*, and *Ecnolagria sp.*, consisted of only short-read Illumina sequence data. Sequences were trimmed using Trimmomatic v0.39 [74] using TruSeq3-PE as reference, and sequences shorter than 25 bp being discarded. The trimmed sequences were assembled using SPAdes v.3.14.1 [75] and binned using Autometa [76]. Sequence data generated from *L. villosa* 2020 and *L. hirta* HG consisted of both short-read Illumina sequence data and long-read Nanopore sequence data, as mentioned above. After trimming, reads were assembled with SPAdes v.3.14.1 as a hybrid assembly with the nanopore flag enabled. Assembled contigs were binned using Autometa [76]. The quality of all MAGs was assessed using CheckM2 v1.0.1 [47] and each MAG was classified using GTDB-Tk v.2.3.2 against database release 214. Coverage reported by SPAdes for each contig was used to calculate the MAG coverage, except for LvStB_2023 where coverage was calculated by read aligned using minimap2 [77].

### Long-read sequencing and assembly of the *L. villosa* 2023 metagenome

The symbiotic organs connected to the reproductive system of six female adults of *L. villosa* were dissected and gently homogenized to release bacterial symbionts. The residual host tissue was separated from the bacterial suspension, and both samples were frozen at -20°C. Later, both samples were thawed and centrifuged for 2 minutes at 3,000 rpm + 2 minutes at 5,000 rpm to pellet the tissue and bacteria, respectively. The supernatant was removed, and 20µL sterile 1x PBS was added to both samples. Genomic DNA was extracted using the Nanobind CBB Big DNA kit (Circulomics, Baltimore, USA) followed by enrichment for HMW (high molecular weight) DNA using the Short Read Eliminator kit XS (Circulomics). Isolated HMW DNA purity and concentrations were measured using Qubit (Thermo Fisher). End-DNA repair and library preparation was performed using the NEBNext Ultra II DNA Library Prep Kit (New England Biolabs, Ipswich, USA) and the Ligation Sequencing Kit V14 (SQK-LSK114; Oxford Nanopore Technologies, Oxford, UK) followed by a clean-up step with AMPure XP beads (Beckman Coulter). Sequencing was performed on a MinION platform (Oxford Nanopore Technologies) and MinION flow cells (vR10.4.1) with 100 ng of the library during a 72 h run.

After sequencing, super-high-accuracy base calling of the raw reads was performed with Guppy v6.3.8 (Oxford Nanopore Technologies) (dna_r10.4.1_400bps_sup.cfg model; split-read function enabled), resulting in a total of 9 Gb sequence data. The resulting reads were *de novo* assembled using Flye v2.9.1 [78, 79] with setting minimum overlap as 10kb and with the “--meta” option, followed by four rounds of polishing with Racon v1.3.3 (Vaser et al., 2017) starting from the Flye assembly with option (-m 8 -x -6 -g -8 -w 500). After each polishing round, reads were re-aligned to the resulting assembly with minimap2 v2.17 [77]. A final round of polishing was performed using Medaka v1.2.0 (https://github.com/nanoporetech/medaka) with the r941_min_high_g344 model using the MinION raw reads. After polishing, haplotype redundancies and overlaps in the assembly based on read depth were purged using Purge_Dups v1.2.6 [80]. The relative contig coverage, GC content and contig taxonomic classification were scanned after each genome assembly using Blobtools and TaxonAnalysis to enable the identification of potential microbial symbiont contigs. We subsequently performed several rounds of Flye assemblies, using only subsets (e.g. 25%) of the complete MinION data and/or read length size-cutoffs (5kb) to optimize symbiont genome assembly.

### Phylogeny of beetles

Mitochondrial genomes (mitogenomes) were recovered from the Eukaryote kingdom bins from each respective sample. Mitogenomes used by Wei *et al.* [28] in their phylogenetic tree were selected for references and outgroups. Mitogenomes from all metagenomic datasets and reference mitogenomes were annotated using the MITOS2 webserver [81] against the RefSe89 Metazoan database, using genetic code 5 (invertebrate mitochondrial). Amino acid sequences of the 13 protein coding genes (PCGs) from each mitogenome were collected and aligned using muscle v5.1 [82]. Nucleic acid sequences of the corresponding PCGs were aligned using pal2nal v14 [83] using the -codontable 5 flag. Nucleic acid alignments were concatenated and a partition file was generated using the pxcat command from the phyx package [84]. Phylogenetic analysis was performed by partitioning each codon position for each gene. AICC model predicted by ModelTest-NG v0.1.7 [85, 86] was used to construct the phylogenetic tree using RAxML-NG v1.2.1 [87] with the parameters --all and --tree pars{25},rand{25}. Alignment file in FASTA format was converted to nexus format using Geneious Prime 2023.2.1 (www.geneious.com). Bayesian analysis was performed by partitioning each codon position for each genes using MrBayes v3.2 [33] with seed and swapseed equal to 42 and using the following parameters lset applyto=(all) nst=6 rates=invgamma; and unlink statefreq=(all) revmat=(all) shape=(all) pinvar=(all); using 10 million generations and sample frequency of 500. The final average standard deviation of split frequencies (ASDSF) was 0.0042.

### *Burkholderia* symbiont phylogeny

Prokka [88] was used to annotate the ORFs of the genomes/ MAGs. Pseudogenes were removed from the MAGs and orthofinder v2.5.5 [89] was run on amino acid sequences of the genomes/ MAGs. A custom script was used to extract the genes with single-copy HOGs that are present in more than 95% (23 HOGs), 90% (126 HOGs), 80% (336 HOGs), 70% (656 HOGs) and 60% (888 HOGs) of the genomes/ MAGs. Muscle v5.1 [82] was used to align the amino acid sequences of the selected HOGs, followed by pal2nal v14 [83] to align the corresponding nucleic acid sequences using -codontable 11. Subsequent steps were similar to those performed while constructing beetle phylogeny. Bayesian analysis was performed using MrBayes v3.2 [33] following the steps and parameters described in beetle mitogenome tree construction. The final ASDSF was 0.0002.

Amino acid sequences of MAGs were blasted (diamond blastP) [90, 91] (with parameters -k 1 -- max-hsps 1 --outfmt 6 qseqid stitle pident evalue qlen slen) against a local copy of the NCBI nr database, where previously identified *Burkholderia* sequences [21, 23] were removed. Genes where the top blastP hits had percent identity less than 50% or those without “Burkholderia” in the subject sequence title of the top hit were classified as putative horizontally transferred genes. These genes along with any pseudogenes were removed from MAGs and orthofinder was used to detect HOGs present in more than 95% (16 HOGs), 90% (98 HOGs), 80% (304 HOGs), 70% (632 HOGs) and 60% (884 HOGs) of the genomes/ MAGs. Subsequent steps were similar to those mentioned in the above paragraph.

### Lagriamide BGC phylogeny

Lagriamide BGC genes from *lgaA* to *lgaI* were extracted. Protein sequences were aligned with muscle v5.1 [82] followed by alignment of DNA sequences using pal2nal v14 [83] using - codontable 11. Pxcat command in the phyx package [84] was used to concatenate the DNA alignments and generate a partition file. Maximum likelihood tree was made using RAxML v8.2.12 (raxmlHPC-PTHREADS-SSE3) [32], with the parameters -f a -# 1000 -p 1989 -x 1989. For the GTRGAMMAI model each gene was partitioned for each codon position, while when using GTRCAT -V model partitioning was only performed per gene as it resulted in higher bootstrap values than partitioning for each codon position in each gene. Bayesian analysis was performed as described in the beetle phylogeny with the final ASDSF being 0.0001.

## Data availability

The data associated with this study was deposited under BioProject accession no. PRJNA1054523. Metagenomic reads have been deposited in the Sequence Read Archive with accessions SRR27332963–SRR27332975. Representative *lga* BGC sequences have been submitted to Genbank with accession numbers PP034267–PP034279. All lagriamide BGC-carrying MAGs were deposited with the following accession numbers: Ecno_1, JAYFRU000000000; Latri_1, JAYFRV000000000; Lgren_6, JAYFRW000000000; LhG_1, JAYFRX000000000; LhHG_2, JAYFRY000000000; LhSB_1, JAYFRZ000000000; Lruf2_2, JAYFSA000000000; Loki_2, JAYFSB000000000; Lruf1_1, JAYFSC000000000; Lv20_9, JAYFSD000000000.

## Acknowledgments

We would like to thank Jean Keller for providing python scripts that helped in creating the symbiont phylogeny. We are grateful to Rebekka Janke, Takema Fukatsu, Kiyoshi Ando, Kimio Masumoto, Juan José López and Rolf Beutel for support with insect collection and to Domenica Schnabelrauch for carrying out PCRs, cloning and Sanger sequencing.

## Author Contributions

SU: Conceptualization, Data curation, Formal Analysis, Investigation, Methodology, Visualization, Writing – original draft, Writing – review & editing.

SCW: Conceptualization, Data curation, Formal Analysis, Investigation, Methodology, Visualization, Writing – original draft, Writing – review & editing.

AN: Data curation, Formal Analysis, Investigation, Methodology HV: Data curation, Formal Analysis, Investigation, Methodology.

LVF: Conceptualization, Investigation, Writing – review & editing, Supervision, Funding acquisition.

MK: Formal Analysis, Methodology, Conceptualization, Investigation, Writing – review & editing, Supervision, Funding acquisition.

JCK: Conceptualization, Investigation, Writing – review & editing, Supervision, Funding acquisition

