## Supplementary Information for "Repeated horizontal acquisition of lagriamide-producing symbionts in Lagriinae beetles"

#### Supplementary figures

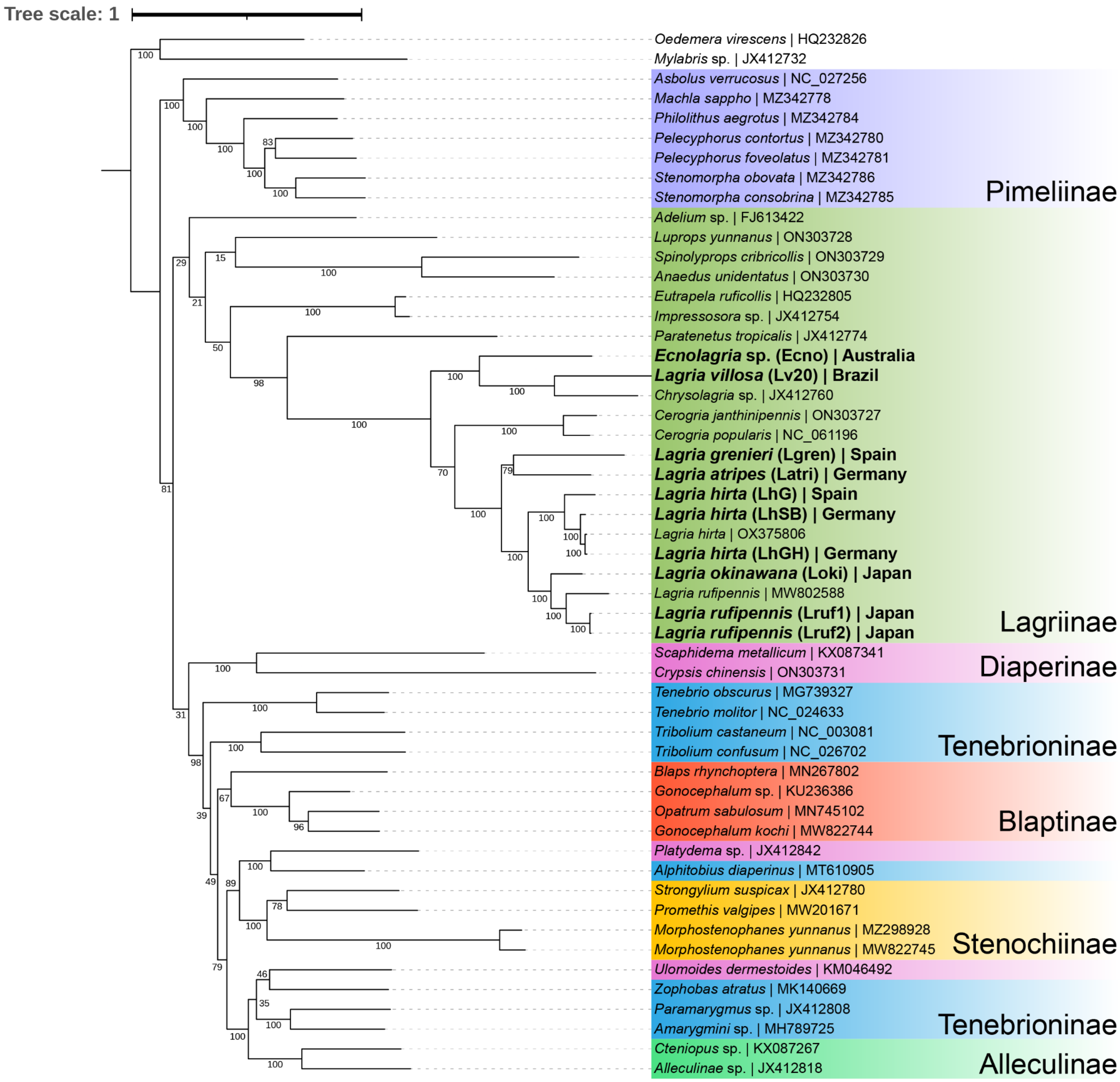

**Fig SI 1**. Beetle mitogenome phylogenetic tree using 13 mitochondrial protein coding genes constructed using RAxML. Branch values represent bootstrap values. Mitogenomes recovered in this study are highlighted with bold lettering.

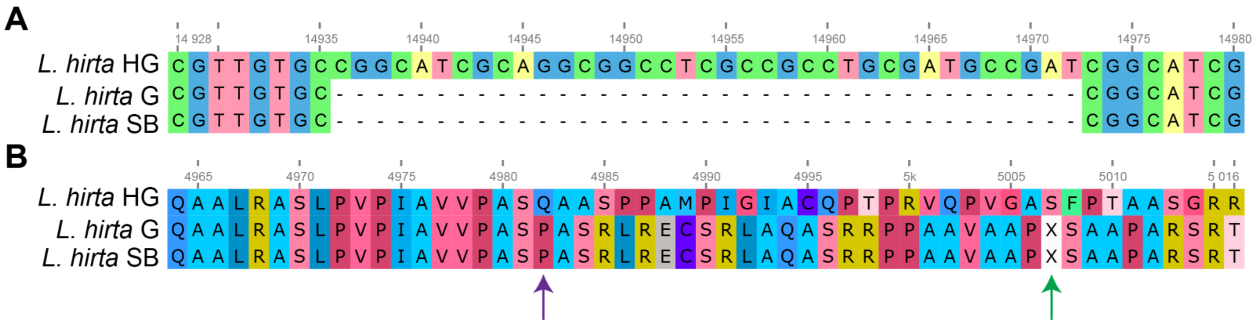

**Fig SI 2**. The break in the *lgaC* gene in the *lga* BGCs from *L. hirta* SB and *L. hirta* G, was found to be a result of A) a 37-bp deletion relative to the complete *lgaC* gene in *L. hirta* HG. This deletion resulted in B) a frameshift (purple arrow) that caused a premature stop codon (green arrow).

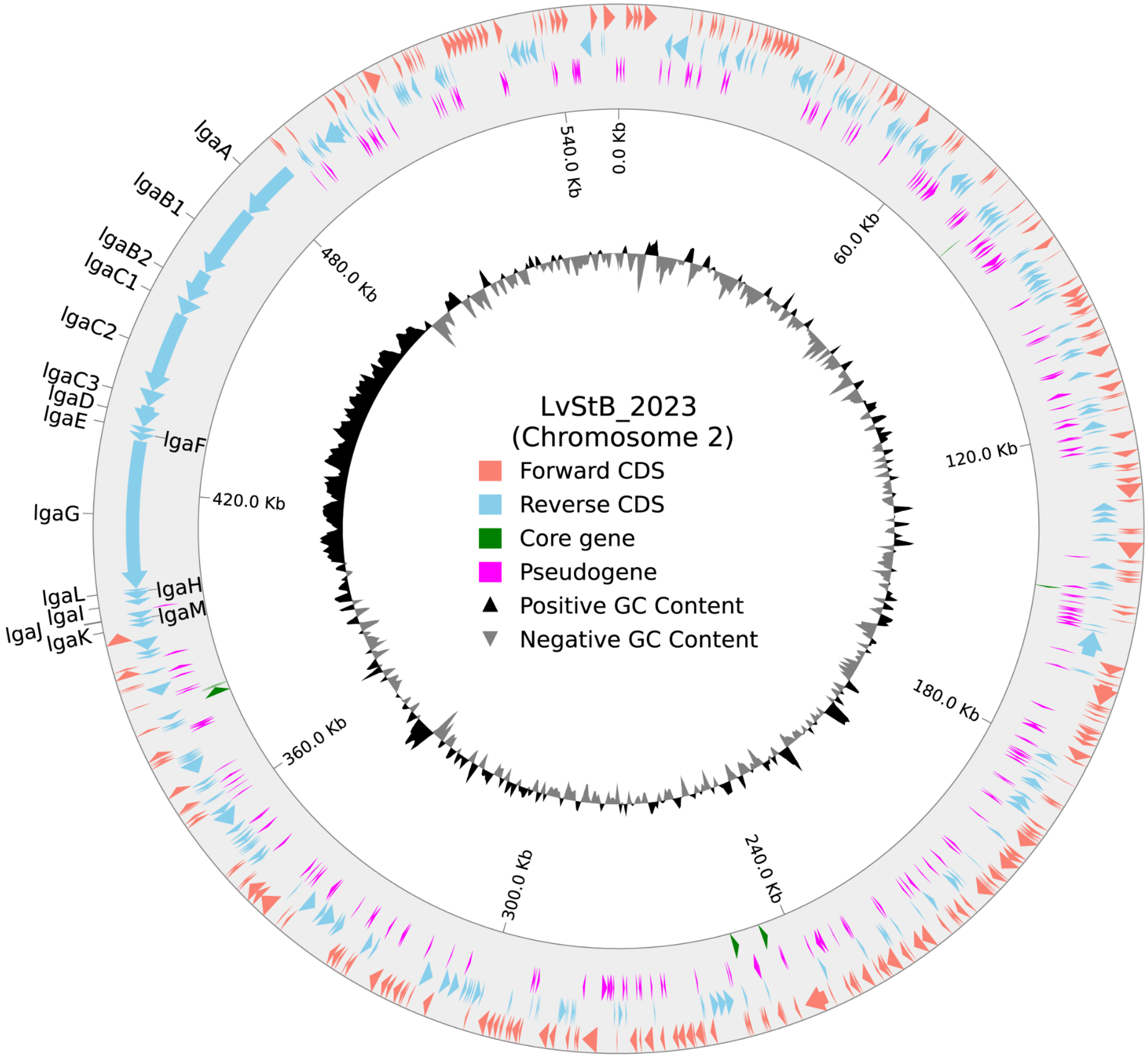

**Fig SI 3**. Circular representation of *lga* BGC harboring chromosome. The increased GC content in the *lga* BGC region provides further evidence towards its horizontal acquisition.

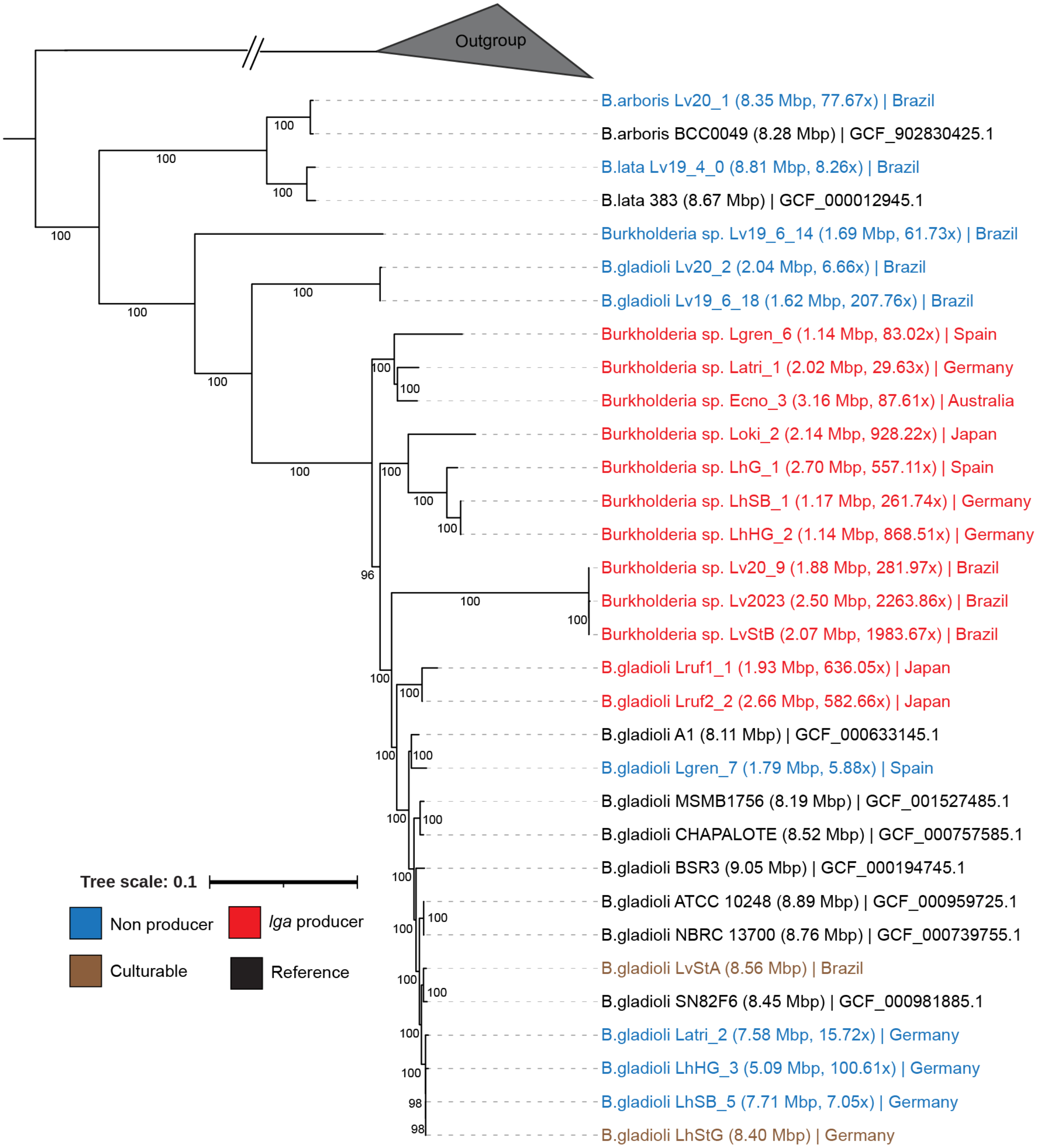

**Fig SI 4.** RAxML phylogenetic tree of Lagriinae beetle associated *Burkholderia* symbionts. Values on nodes represent bootstrap values. Genome size and coverage are represented in brackets next to MAG ID. Outgroups include - *Paraburkholderia acidiphila* (GCF_009789655.1), *Cupriavidus necator* (GCF_000219215.1), *Herbaspirillum seropedicae* (GCF_001040945.1).

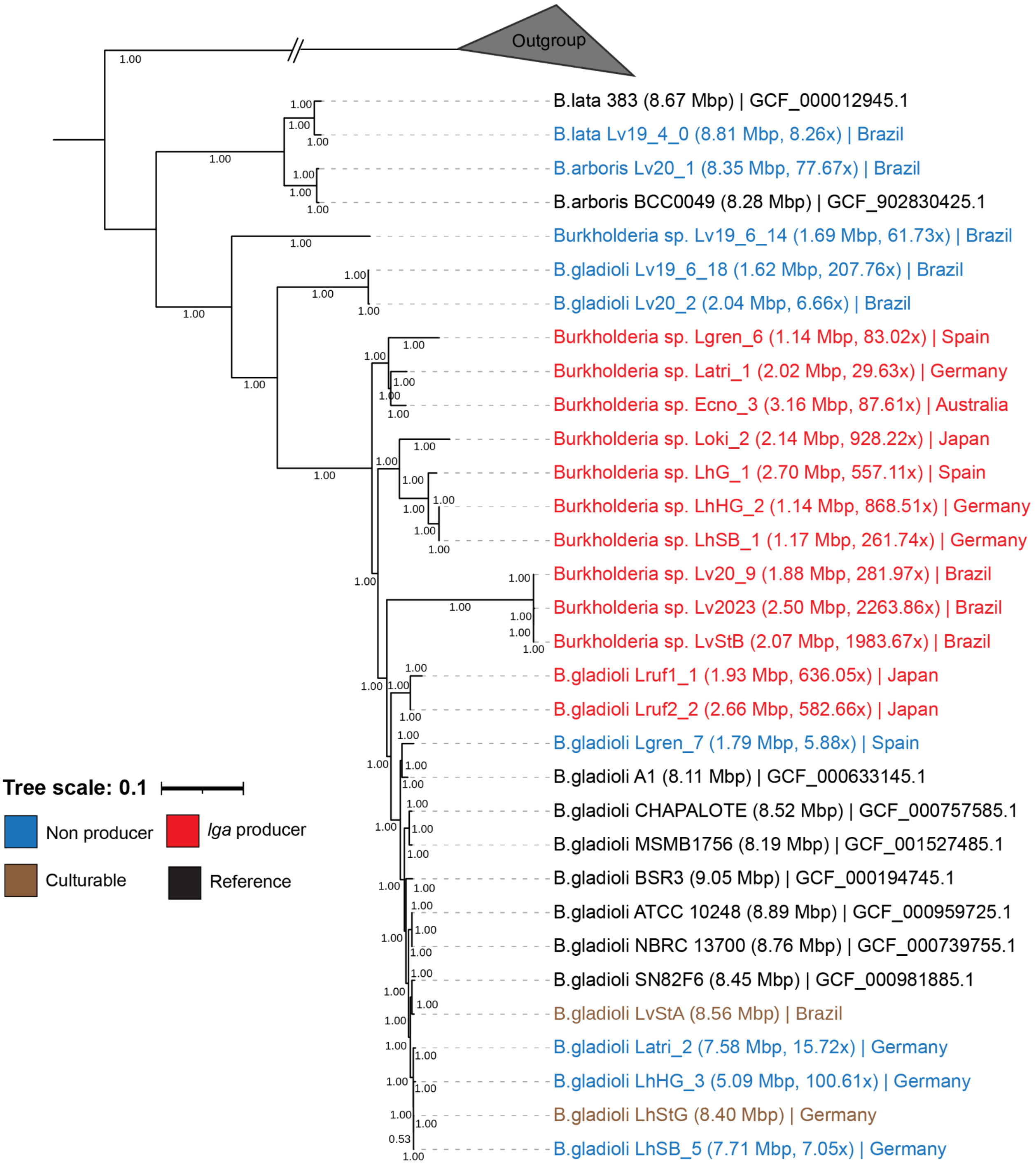

**Fig SI 5.** Bayesian phylogenetic tree of Lagriinae beetle associated *Burkholderia* symbionts. Values on nodes represent posterior probabilities. Genome size and coverage are represented in brackets next to MAG ID. Outgroups include - *Paraburkholderia acidiphila* (GCF_009789655.1), *Cupriavidus necator* (GCF_000219215.1), *Herbaspirillum seropedicae* (GCF_001040945.1).

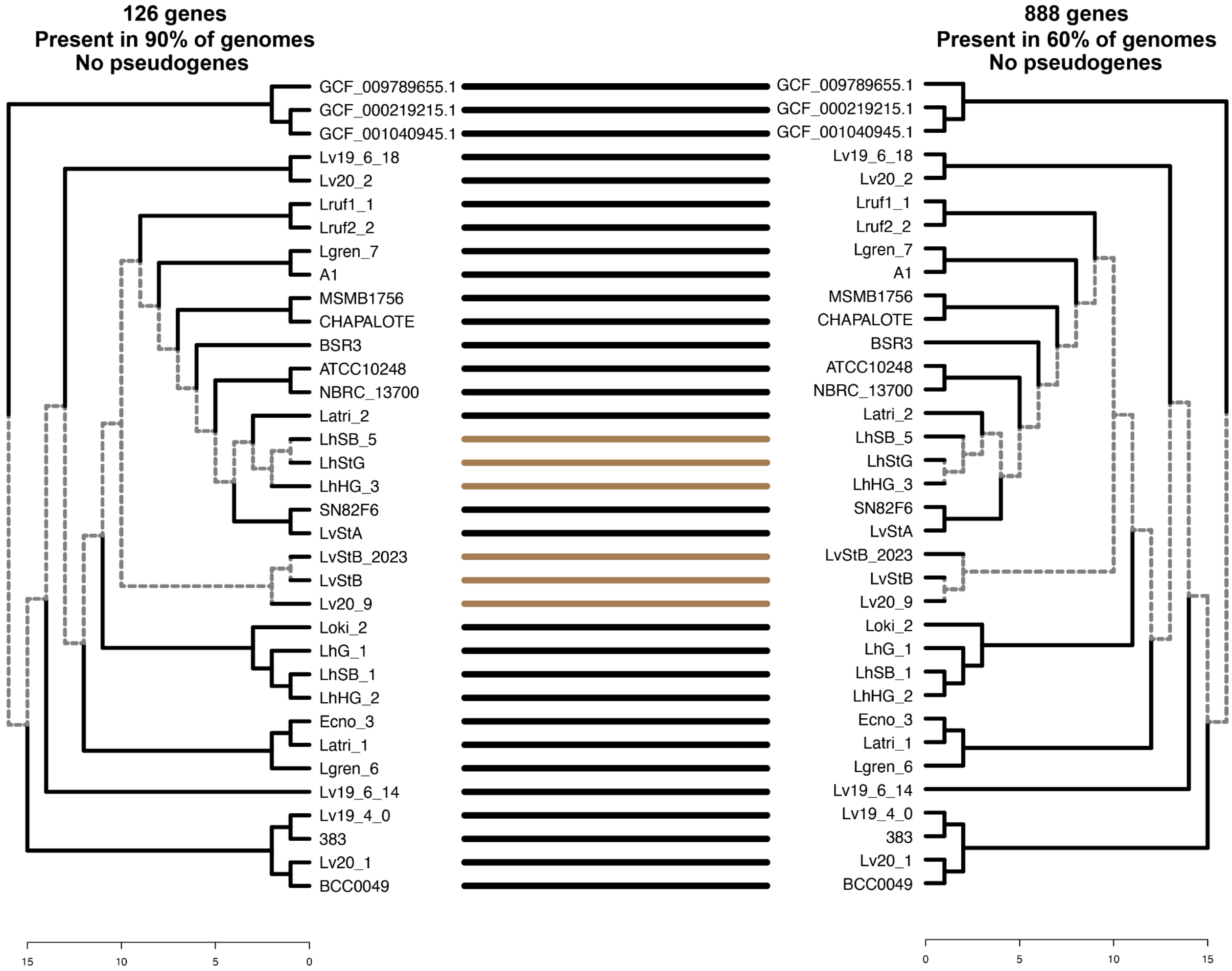

**Fig SI 6.** Tanglegram between *Burkholderia* symbiont phylogeny constructed using a different number of genes. Both phylogenies show a non-monophyletic clade of lagriamide-containing *Burkholderia* symbionts, and are highly similar other than very minor discrepancies in the terminal nodes. Node labels represent MAG or strain ID. GCF_009789655.1 (*Paraburkholderia acidiphila*), GCF_000219215.1 (*Cupriavidus necator*), and GCF_001040945.1 (*Herbaspirillum seropedicae*) represent outgroups used. Dashed lines represent nodes that are unique between the respective phylogenies.

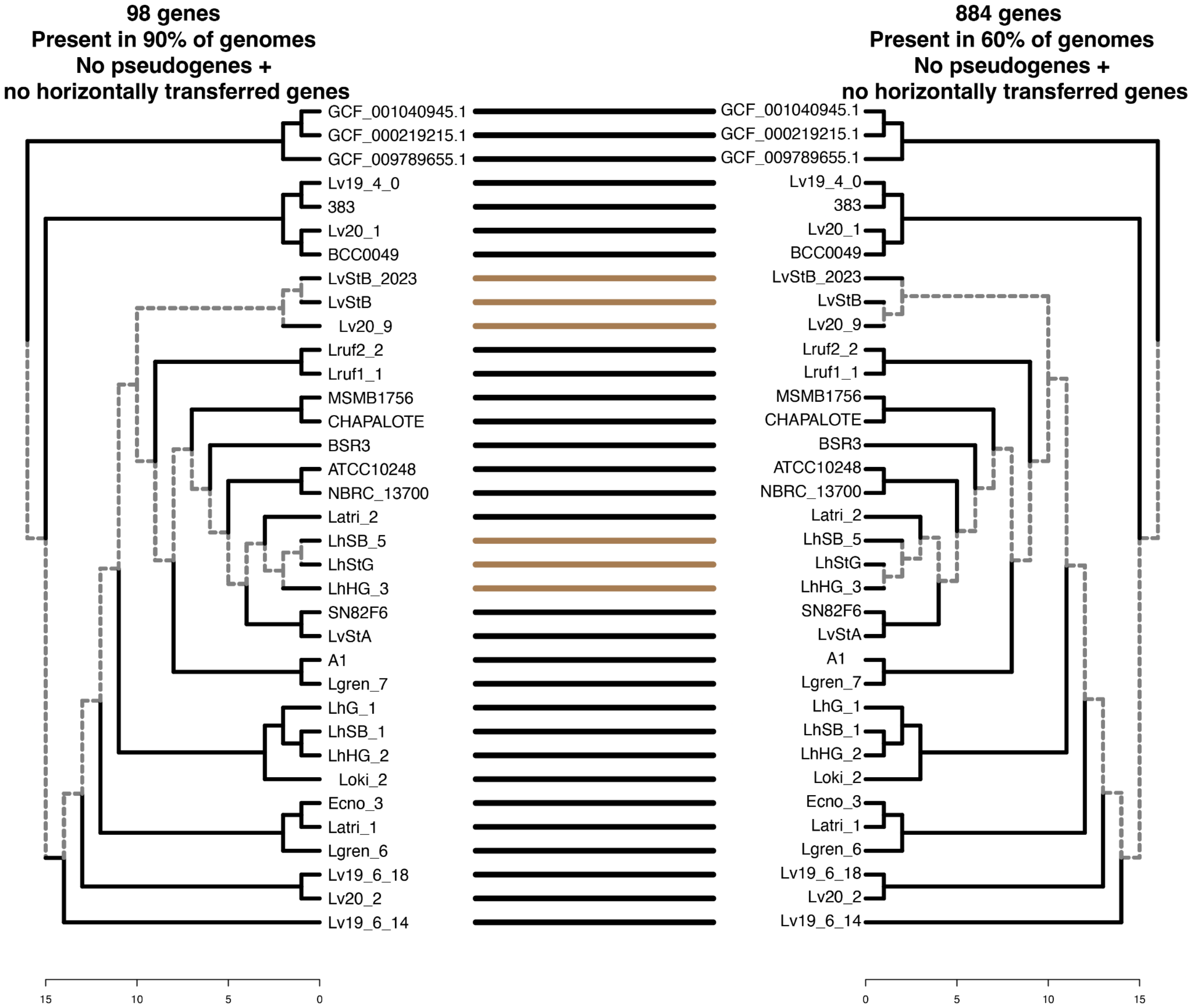

**Fig SI 7.** Tanglegram between *Burkholderia* symbiont phylogeny constructed using different number of genes. Both phylogenies show a non-monophyletic clade of lagriamide-containing *Burkholderia* symbionts, and are highly similar other than very minor discrepancies in the terminal nodes. Node labels represent MAG or strain ID. GCF_009789655.1 (*Paraburkholderia acidiphila*), GCF_000219215.1 (*Cupriavidus necator*), and GCF_001040945.1 (*Herbaspirillum seropedicae*) represent outgroups used. Dashed lines represent nodes that are unique between the respective phylogenies.

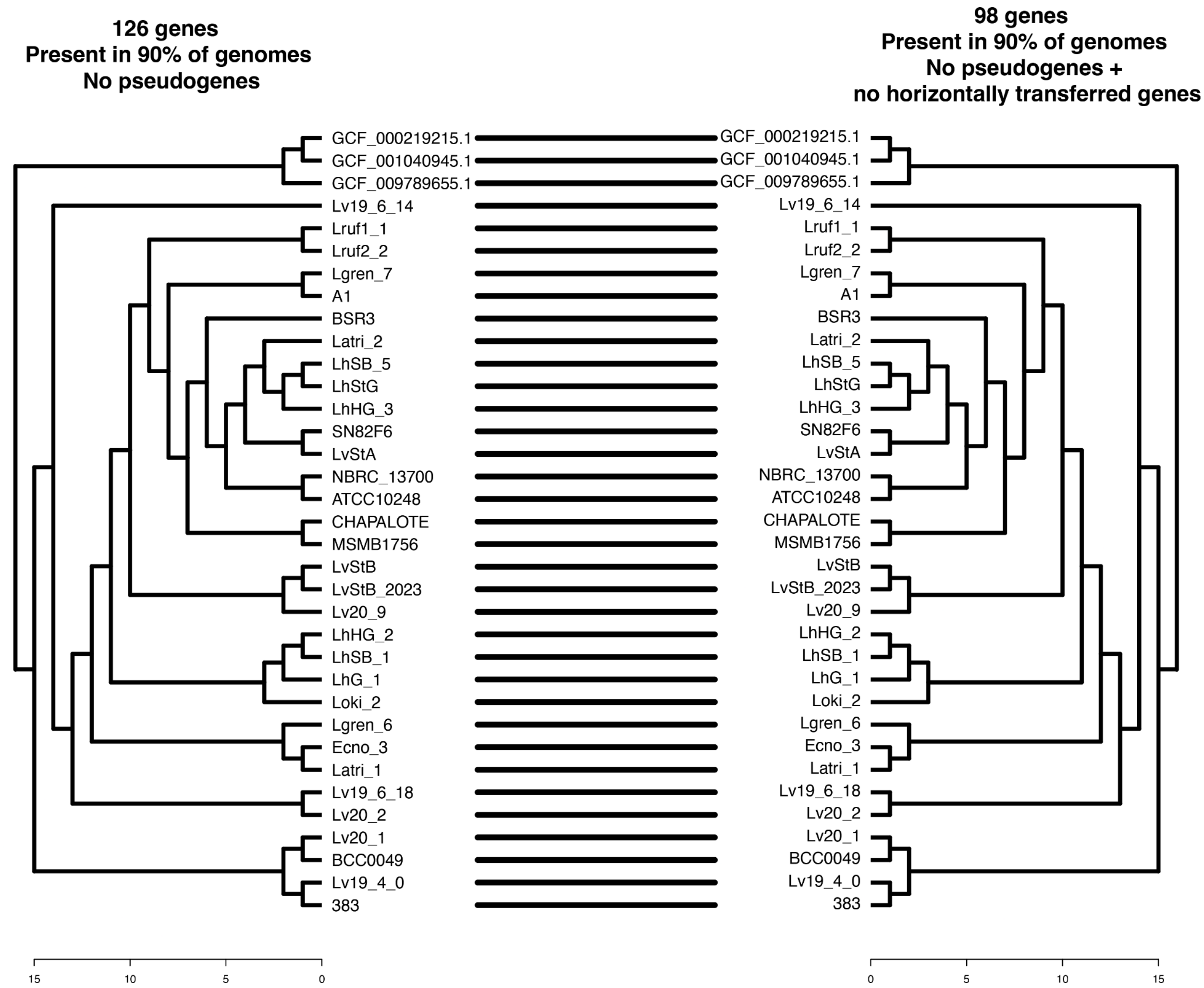

**Fig SI 8.** Tanglegram between *Burkholderia* symbiont phylogeny constructed using different number and types of genes. Both phylogenies show a non-monophyletic clade of lagriamide-containing *Burkholderia* symbionts, and are exactly the same. Node labels represent MAG or strain ID. GCF_009789655.1 (*Paraburkholderia acidiphila*), GCF_000219215.1 (*Cupriavidus necator*), and GCF_001040945.1 (*Herbaspirillum seropedicae*) represent outgroups used.

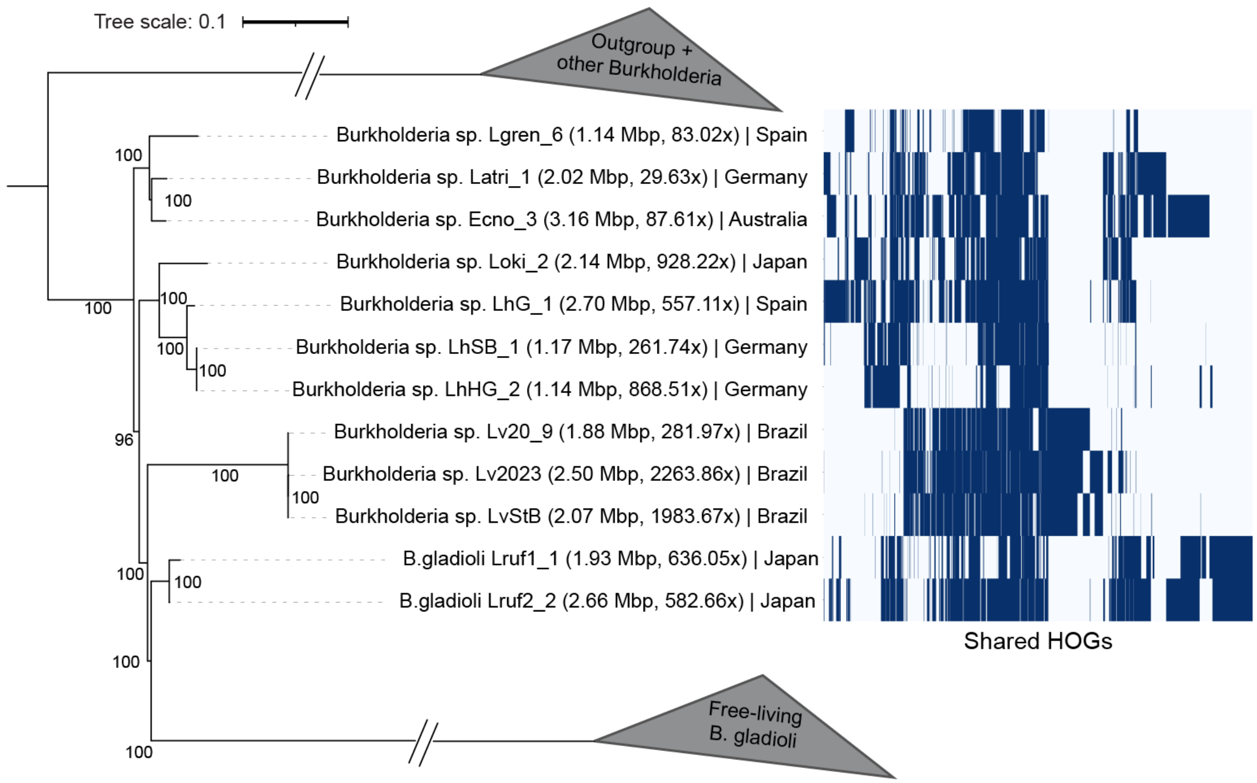

**Fig SI 9**. RAxML phylogenetic tree (left) and shared hierarchical orthogroups (HOGs) (non-pseudogenes) between different *lga* BGC carrying *Burkholderia* symbionts (matrix on the right). Each blue line indicates a shared HOG. HOGs have been hierarchically clustered on the x-axis to improve visualization. Values on nodes indicate bootstrap values. Genome size and coverage is represented in brackets next to MAG ID.

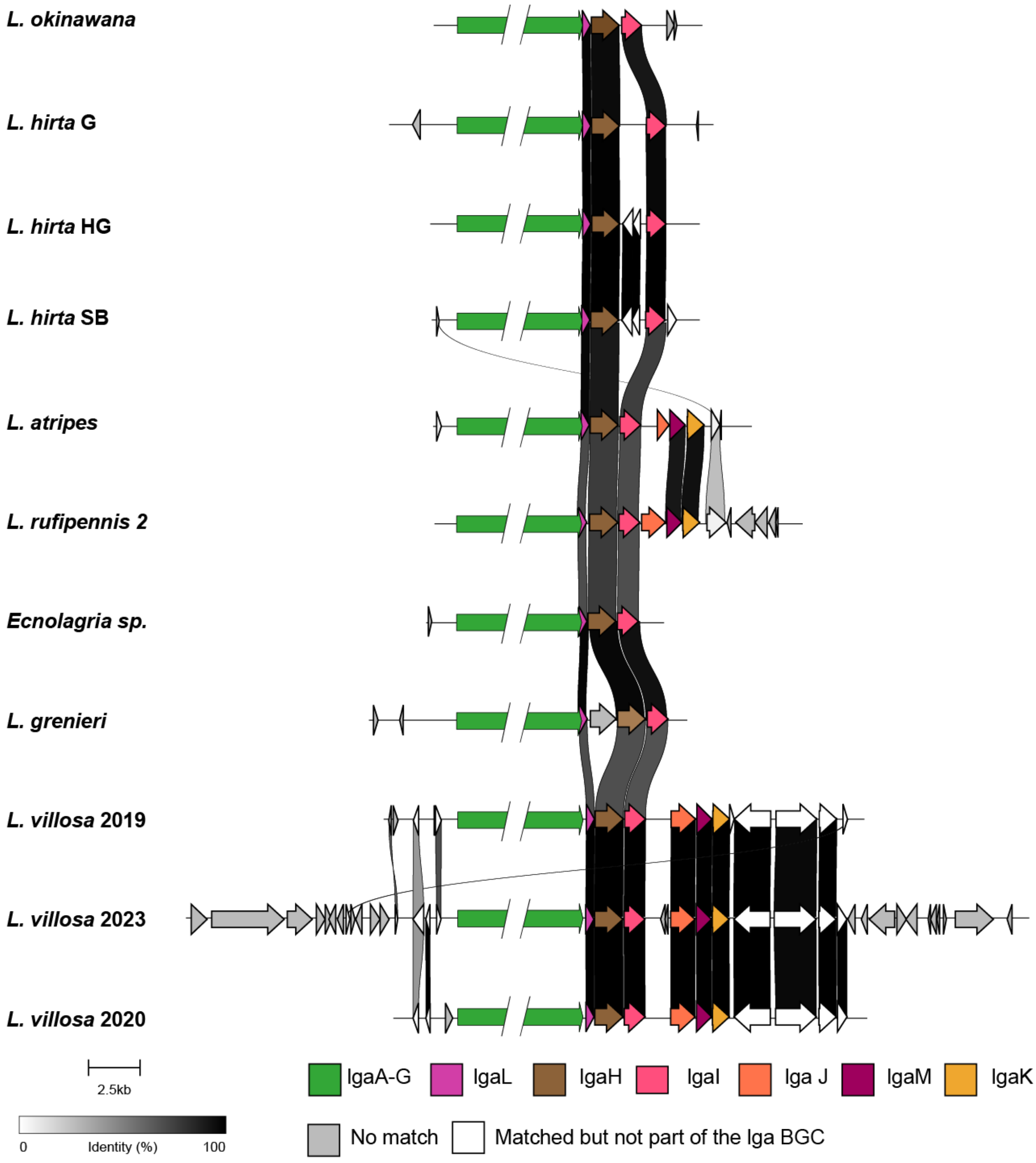

**Fig SI 10**. Comparison of the gene flanking the *lga* BGCs. The lack of similarity between the flanking genes across different BGCs indicates a lack of synteny.

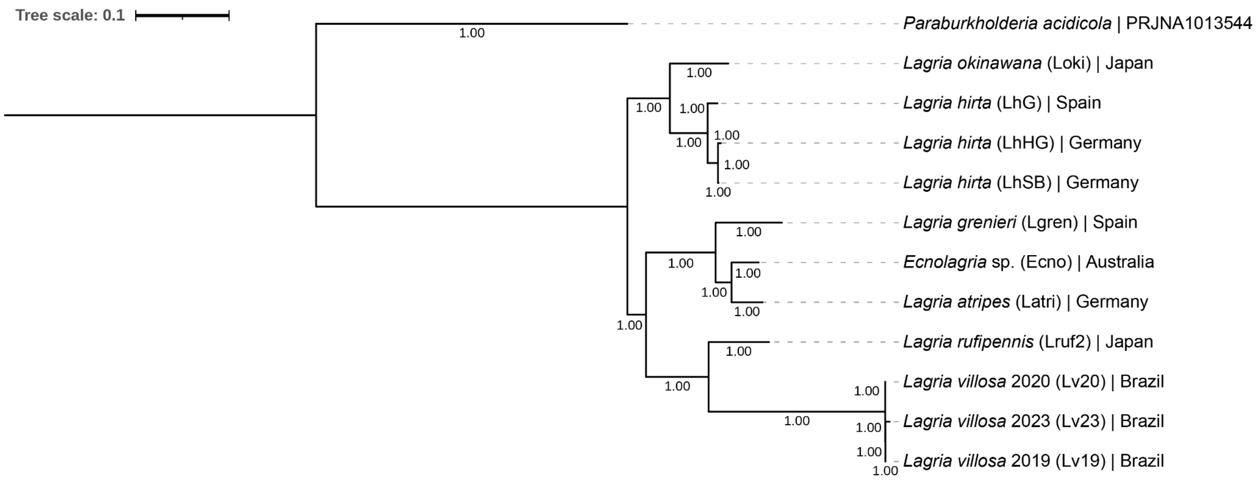

**Fig SI 11**. Bayesian phylogeny of the *lga* BGCs. Values on nodes represent posterior probabilities.

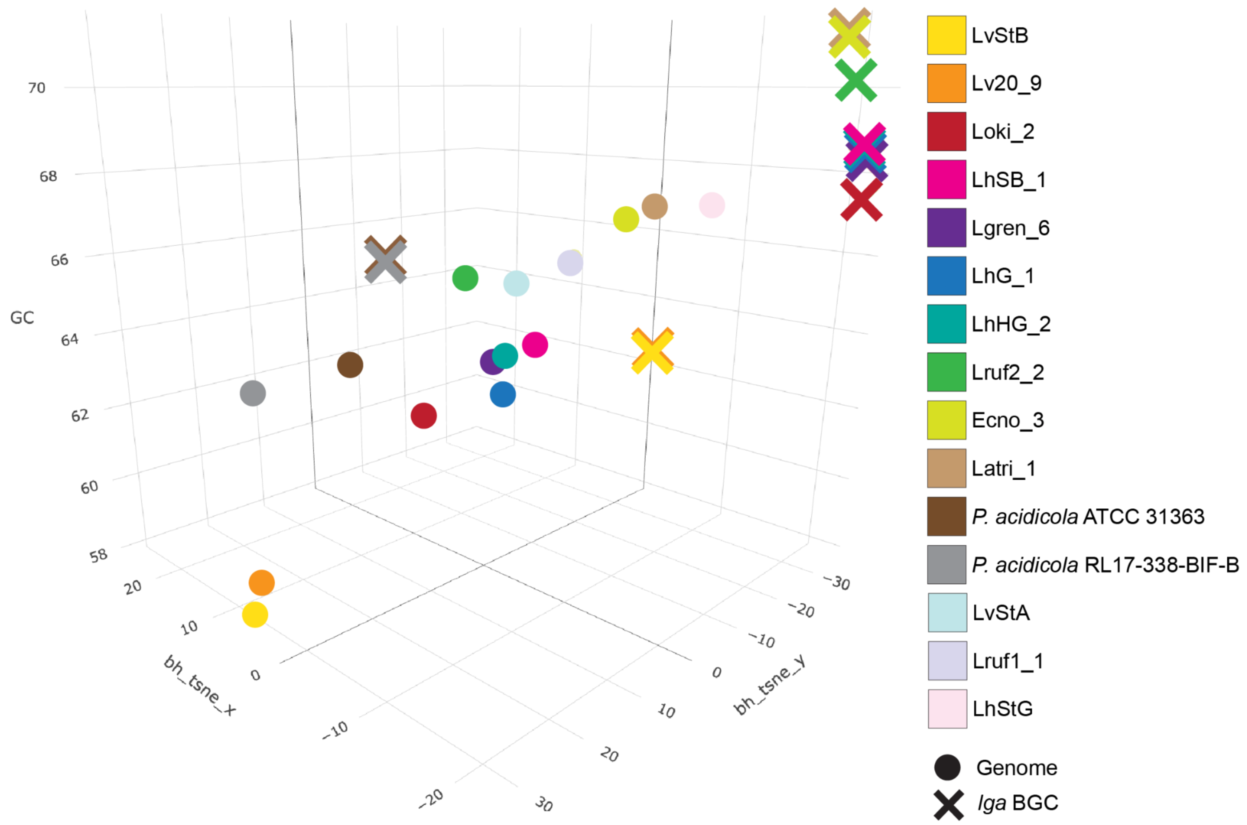

**Fig SI 12.** Dimension reduction of 5-mer nucleotide composition (x,y-axes) of lagriamide BGCs and *Burkholderia* genomes relative to GC content (z-axis).

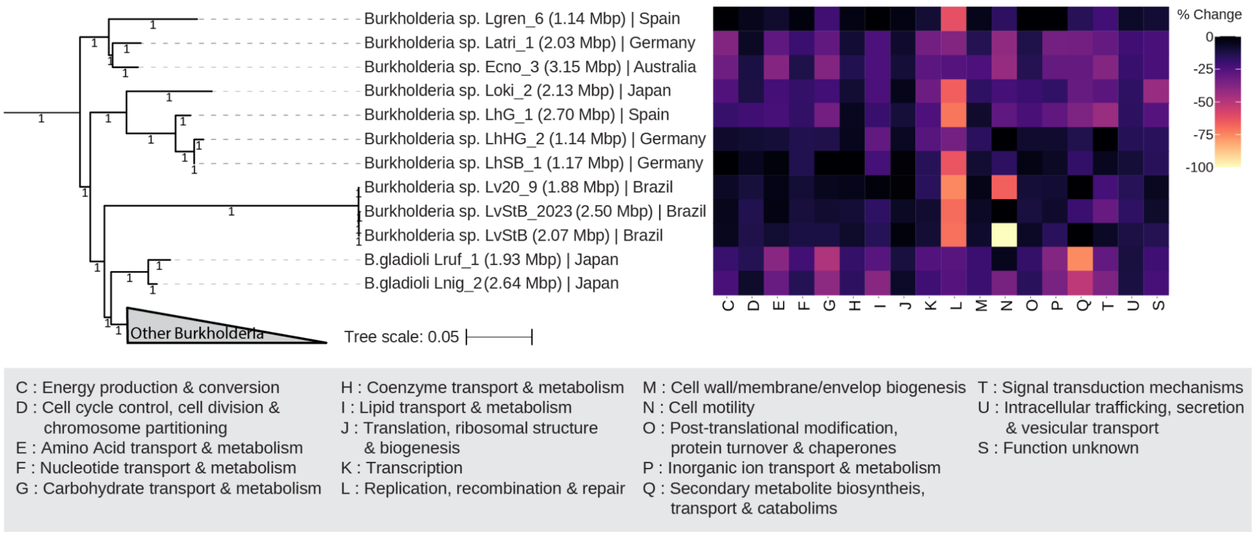

**Fig SI 13.** The percentage change in COG-annotated genes between gene datasets with and without pseudogenes in each of the beetle-associated, *lga*-carrying *Burkholderia* symbionts.

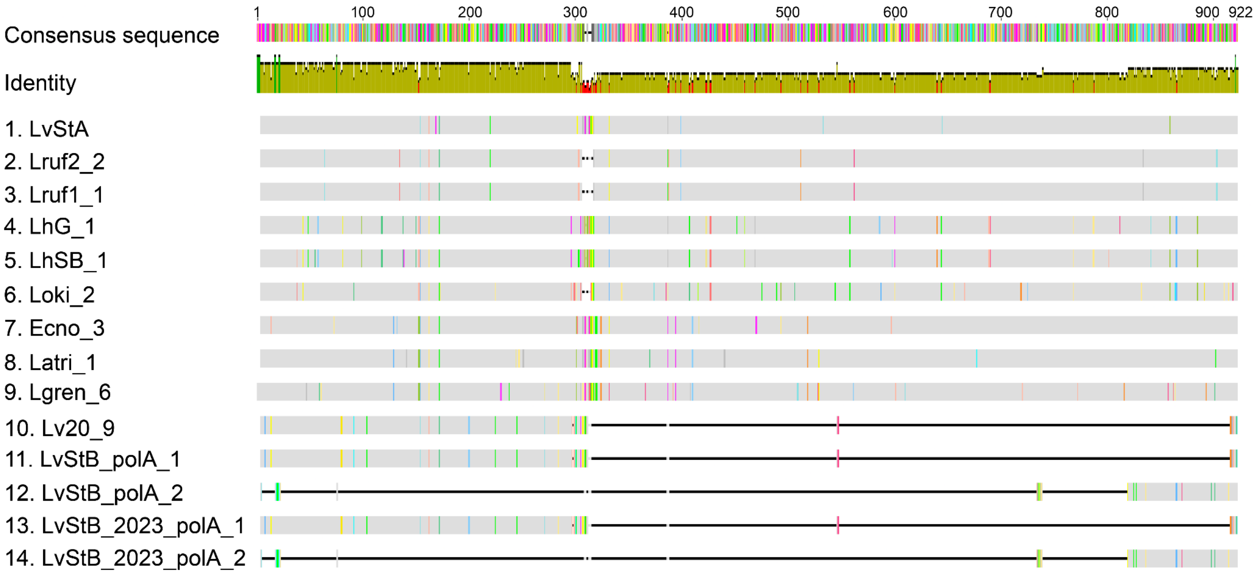

**Fig SI 14.** Multiple sequence alignment of DNA polymerase I gene (*polA*) in different lagriamide-carrying symbionts. LvStA is used as a reference. Colored blocks within the alignment indicate disagreements with the consensus sequence.

### Supplemental methods

#### Barcoding of *Lagria rufipennis* and *Lagria nigricollis* specimens

For barcoding, 19 *Lagria rufipennis* and 10 *Lagria nigricollis* specimens (based on morphological identification) were available. One leg of each specimen was used for DNA extraction using the Epicenter MasterPure extraction kit. A part of the cytochrome oxidase I (COI) gene was amplified using the following primer combinations (**Table SI 1**): CLepFolF + CLepFolR; LCO1490 + HCO2198; dgLCO1490 + dgHCO2198; C1-J-2183-F + C1-N-2609-R. PCRs were carried out with initial denaturation at 94°C for 3 minutes, first 10 cycles of denaturation at 94°C for 40 seconds, annealing at 55°C for 40 seconds and elongation at 72°C for 60 seconds followed by 35 cycles of denaturation at 94°C for 40 seconds, annealing at 45°C for 40 seconds and elongation at 72°C for 60 seconds, final elongation step at 72°C for 5 minutes. For the last three primer combinations the first 10 cycles were omitted and annealing temperatures adjusted to 42°C (LCO1490 + HCO2198; dgLCO1490 + dgHCO2198) and 48°C (C1-J-2183-F + C1-N-2609-R). Successfully amplified products were purified using the Zymo Research DNA Cleanup & Concentrator Kit according to manufactures instructions and Sanger sequenced using forward and reverse primers. Consensus Sequences were created and aligned using Geneious Prime. For assignment to the two different species, a phylogenetic tree including all barcode sequences and the COI sequences extracted from the Lruf1 and Lruf2 metagenomes as well as a L. rufipennis sequence available on NCBI (MW802588) was reconstructed using FastTree 2 [1].

**Table SI 1**: Primers used for COI barcoding of *L. rufipennis* and *L. nigricollis* specimens.

| PCR# | Name | Fw/Rv | 5' -> 3' sequence | Product size | Target | Reference |
|---|---|---|---|---|---|---|
| 1 | CLepFolF (LepF1) | Fw | ATTCAACCAATCATAAAGATATTGG | ~700 bp | Insects and other eukaryotic CO I | [2, 3], |
| | CLepFolR (LepR1) | Rv | TAAACTTCTGGATGTCCAAAAAATCA | | | |
| 2 | LCO1490 | Fw | GGTCAACAAATCATAAAGATATTGG | | | [4] |
| | HCO2198 | Rv | TAAACTTCAGGGTGACCAAAAAATCA | | | |
| 3 | dgLCO1490 | Fw | GGTCAACAAATCATAAAGAYATYGG | | | [5] |

| PCR# | Name | Fw/Rv | 5’ -> 3’ sequence | Product size | Target | Reference |
|---|---|---|---|---|---|---|
| | dgHCO2198 | Rv | TAAACTTCAGGGTGACCAAARAAYCA | | | |
| 4 | C1-J-2183 | Fw | CAACATTTATTTTGATTTTTTGG | ~500 bp | Pyrrhocoridae COX1 | [6] |

#### K-mer analysis of BGCs and associated genomes

The contigs of all *Burkholderia* bins, including their BGCs, were concatenated into a single fasta file and the autometa-kmers method was implemented to find the 5-mer frequency distribution patterns of each contig. The norm-method parameter was set to “am-clr”, the pca-dimensions parameter set to 50, the embedding-method parameter was set to “bhsne”, with 2 embedding dimensions, and a seed set at 42. This resulted in a set of x,y coordinates for each contig representing the dimension-reduced position of its kmer frequency. The GC content of the contig was added as a third dimension.

Visualization of k-mer frequencies of the BGCs and the genomes revealed three clusters of BGCs: The BGCs from the two soil-borne *Paraburkholderias*, the BGCs from the two Brazilian *L. villosa*-derived LvStB strains, and then all others (**Fig. SI 7**). However, a similar pattern can be observed for the nucleotide composition of the respective genomes wherein LvStB and Lv20_9 are distant from all others, as are the points representing the two soil-borne *Paraburkholderias*. All other lga-carrying *Burkholderia* and cultured *Lagria*-associated genomes (LvStA and LhStG) form a third loose cluster. None of the BGCs share similar k-mer composition with their respective genomes suggesting that all are likely horizontally acquired from an independent donor organism.

#### COG annotation of beetle-associated *lga*-carrying *Burkholderia* symbiont genomes

All genomes for all beetle-associated *lga*-carrying *Burkholderia* symbionts were annotated against the COG database using eggNOG-mapper v2.1.9 [7, 8]. Input used included one dataset of all genes identified using prokka [9], and a second dataset where identified pseudogenes had been removed. The number of genes assigned per COG category was counted and the percentage change between the datasets with and without pseudogenes was calculated for each dataset pair.

Here, we noted that Category L (DNA replication, recombination, and repair) had the highest number of pseudogenes in many of the MAGs, with the highest counts in the three *L. villosa*-associated *Burkholderia* (**Fig. SI 8**). We also noted a number of genes assigned within Category N (Cell motility) appeared to be pseudogenes in two of the three *L. villosa*-associated MAGs.

#### General bioinformatic analyses

Phylogenetic trees were visualised and modified in the Interactive Tree of Life server [10]. Tanglegrams were made using the dendextend [11] package in RStudio [12]. The circular genome of LvStB_2023 was made using pyCirclize (https://github.com/moshi4/pyCirclize) package in Jupyter Notebook. COG categories were identified using eggNOG-mapper v2.1.9 [7, 8]. For identification of pseudogenes, diamond blastP [13, 14] alignment was performed for amino acid sequences of ORFs against a local copy of the NCBI nr database (with parameters -k 1 --max-hsps 1 --outfmt 6 qseqid stitle pident evalue qlen slen). Previously published *Burkholderia* sequences [15, 16] were removed from the local nr database. ORFs with a length less than 80% of their closest blastP hit were classified as pseudogenes, as described previously [16]. Transposases were identified by parsing Prokka [9] and DRAM v1.4.6 (using --use_uniref flag) [17, 18] output for "transposase". Set of core genes observed to be present in most reduced symbionts were taken from Table 2 of McCutcheon and Moran [19]. PlasFlow [20], PLASMe (https://github.com/HubertTang/PLASMe), and plasmidVerify (https://github.com/ablab/plasmidVerify) were used to verify the plasmid assignment of the smallest contig in the LvStB_2023 genome. ANI and AF analysis was done using skani v0.2.1 [21].

#### Heatmap of shared HOGs

Pseudogenes were removed from the MAGs and Orthofinder v2.5.5 [22] was run on relevant genomes. Shared HOGs (single and multiple copy) were converted into a presence/absence matrix for the heatmap. Hierarchical clustering of the HOGs (x-axis in **Fig. 4** and **Fig. SI 9**) was done using the jaccard distance and ward method in Python.

#### Confirmation that BGC contigs were binned correctly

Each of the contigs with the recovered *lga* BGCs had been assigned to a genomic bin during the binning process. To validate that this binning was correct we first assessed whether the coverage and nucleotide composition of the contig(s) carrying the *lga* BGC approximately matched that of the bin to which it was assigned. We then performed paired-end connection mapping of reads mapped back to the assembled contigs with BBMap (https://sourceforge.net/projects/bbmap), for each metagenomic sample, using the cytoscapeviz.pl script from the Multi-metagenome package [23]. We determined which bins the contigs with the greatest number of connections to the *lga* BGC contig were assigned and compared whether this was in agreement with the assigned bin of the BGC. According to these two criteria, it appeared that the contig(s) carrying the *lga* BGC had been binned correctly. This process is described in detail below.

In all cases of lagriamide BGC recovery, we followed the same protocol to determine if a contig carrying a predicted lagriamide BGC had been binned correctly:

1. Visual inspection of the k-mer frequency and coverage of the BGC carrying coverage relative to bins. This was achieved using the tables of dimension-reduced representation of contig 5-mer frequencies generated by Autometa [24] as input in the following R code which employed plotly:

```
library(plotly)

data <- read.table("sample.txt", header = TRUE, sep = "\t")

#3D scatter plot, points scaled by length of contig

p <- plot_ly(data, x = ~x, y = ~y, z = ~coverage, color = ~bin, text =
~paste('Contig:',contig,'<br>Bin:', bin)) %>%
  add_markers() %>%
  layout(scene = list(xaxis = list(title = 'bh_tsne_x'),
                   yaxis = list(title = 'bh_tsne_y'),
                   zaxis = list(title = 'cov')))
```

   All input data is available upon request. Please .

2. We performed paired-end connection mapping of reads mapped back to the assembled contigs with BBMap (https://sourceforge.net/projects/bbmap/), for each metagenomic sample, using the cytoscapeviz.pl script from the Multi-metagenome package [23]. We determined which bins the contigs with the greatest number of connections to the *lga* BGC contig were assigned and compared whether this was in agreement with the assigned bin of the BGC.

#### *L. villosa* 2020

Binning via Autometa [24] placed the contig predicted to carry the lagriamide BGC, contig NODE_78_length_99421_cov_185.181894 in bin Lv20_9. Visualization of the dimension-reduced embedded 5-mer frequency data revealed that the contig was found at approximately the same abundance (coverage) as other contigs present in the Lv20_9, but exhibited a divergent 5-mer frequency pattern from the other contigs in Lv20_9 (Fig. M1). This divergence is likely a result of the horizontal transfer of the BGC as observed for the *lga* BGC in other LvStB bins associated with *L. villosa* beetles.

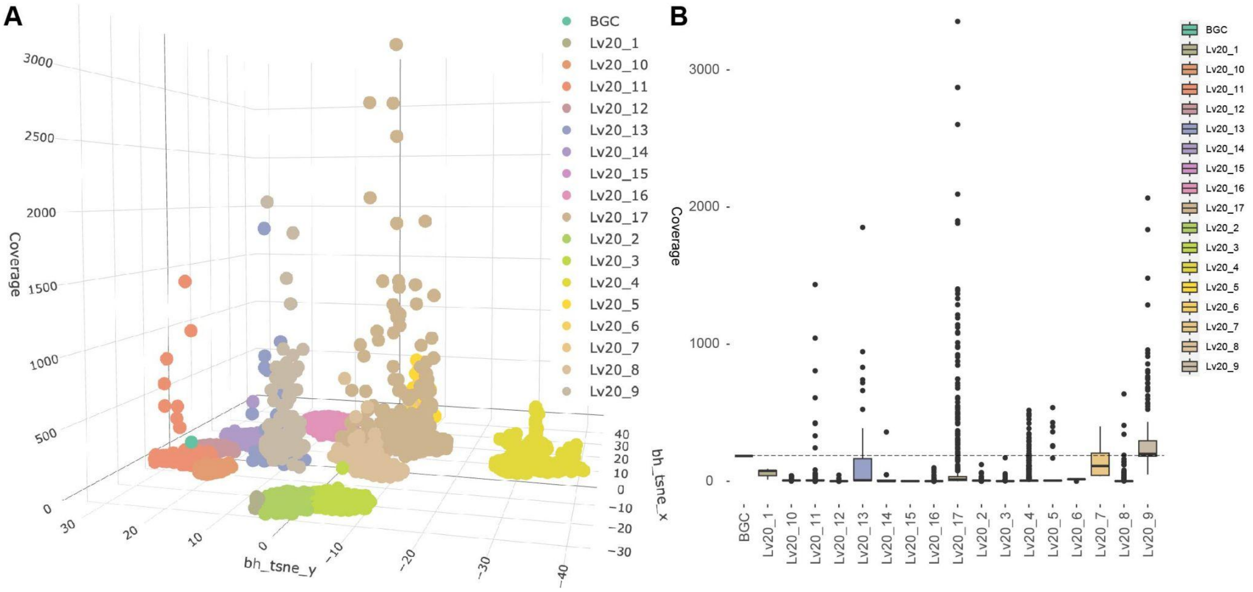

**Figure M1.** Placement of BGC contigs relative to MAGs. A) Three-dimensional visualization of all binned bacterial contigs from the *L. villosa* 2020 metagenome. Each point represents an assembled contig colored by the bin in which it was clustered. The contigs on which the lagriamide BGC was identified have been indicated. B) Range of contig coverages (a proxy for abundance) of each MAG and the lga BGC contig(s). A dotted line is provided for easy comparison to find MAG(s) that have coverages within the range of the BGC coverage.

Reads were mapped back to all scaffolds and connections between contigs were inferred and counted using pair-end mapping. During the binning process, all contigs smaller than 3000 bp were discarded. Therefore any connected contigs with a length less than 3000 bp could not be used to determine if the clustering of the BGC contig (NODE_78_length_99421_cov_185.181894) in bin Lv20_9 was correct. All binned contigs that were predicted as connected to the BGC contig were in bin Lv20_9 (Table M1) and we therefore could confidently conclude that the BGC contig in this sample was binned correctly. However, as the BGC contig was predicted to be connected to several contigs within this MAG, we cannot discount the possibility that the BGC is present in different loci following genome rearrangements within closely related strains represented by this MAG.

**Table M1.** Number of predicted connections shared between the contig with the Iga BGC and other binned contigs in the *L. villosa* 2020 sample

| BGC contig | Connected contig | # connections | Connected contig bin |
|---|---|---|---|
| NODE_78_length_99421_cov_185.181894 | NODE_866_length_15573_cov_240.468471 | 110 | Lv20_9 |
| NODE_78_length_99421_cov_185.181894 | NODE_3563_length_5833_cov_265.473887 | 92 | Lv20_9 |
| NODE_78_length_99421_cov_185.181894 | NODE_1005_length_14019_cov_261.557515 | 57 | Lv20_9 |
| NODE_78_length_99421_cov_185.181894 | NODE_431_length_25132_cov_182.634193 | 45 | Lv20_9 |
| NODE_78_length_99421_cov_185.181894 | NODE_567_length_21169_cov_208.198935 | 28 | Lv20_9 |
| NODE_78_length_99421_cov_185.181894 | NODE_2341_length_7858_cov_280.177726 | 24 | Lv20_9 |
| NODE_78_length_99421_cov_185.181894 | NODE_1655_length_10067_cov_197.738129 | 23 | Lv20_9 |
| NODE_78_length_99421_cov_185.181894 | NODE_895_length_15245_cov_379.362416 | 21 | Lv20_9 |
| NODE_78_length_99421_cov_185.181894 | NODE_5571_length_4157_cov_664.858809 | 12 | Lv20_9 |
| NODE_78_length_99421_cov_185.181894 | NODE_5802_length_4036_cov_2066.283960 | 11 | Lv20_9 |
| NODE_78_length_99421_cov_185.181894 | NODE_2447_length_7602_cov_226.291505 | 6 | Lv20_9 |

#### *L. okinawana*

Binning via Autometa placed the contigs predicted to carry the lagriamide BGC, contigs NODE_46_length_35697_cov_543.659124, NODE_161_length_14264_cov_611.590118, NODE_180_length_12254_cov_531.844379, NODE_311_length_7428_cov_485.445246, NODE_348_length_6767_cov_572.22242 in bin Loki_2. Visualization of the dimension-reduced embedded 5-mer frequency data revealed that the contig was found at approximately the same abundance (coverage) as other contigs present in the Loki_2, and shared the same approximate 5-mer frequency pattern (Fig. M2)

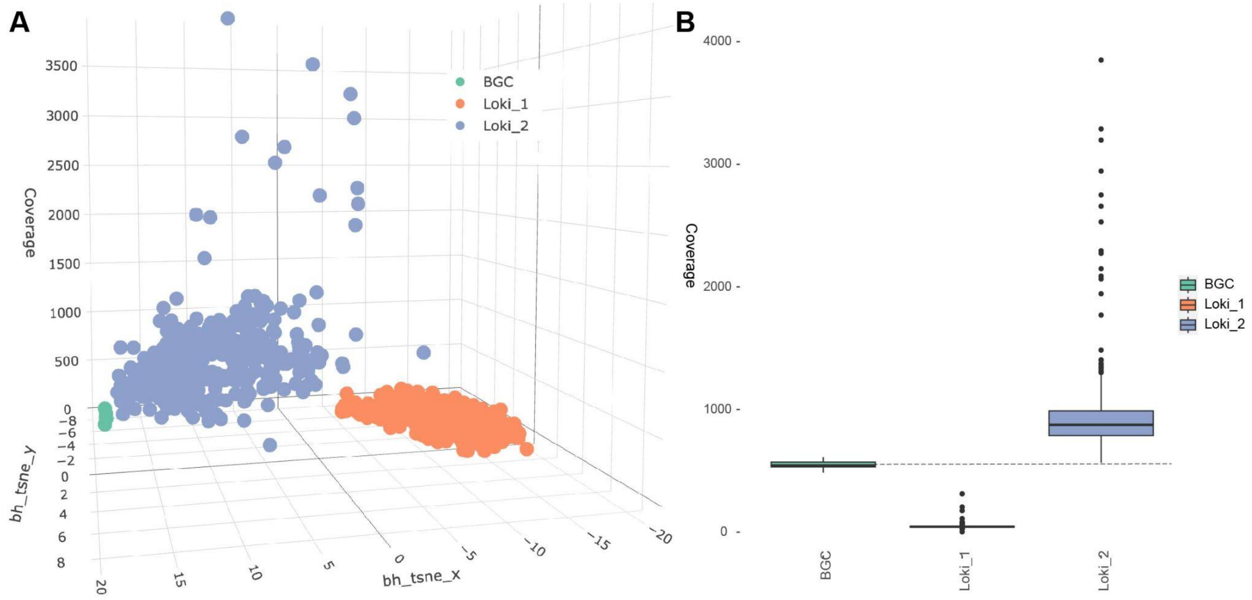

**Figure M2.** Placement of BGC contigs relative to MAGs. A) Three dimensional visualization of all binned bacterial contigs from the *L. okinawana* metagenome. Each point represents an assembled contig colored by the bin in which it was clustered. The contigs on which the lagriamide BGC was identified have been indicated. B) Range of contig coverages (a proxy for abundance) of each MAG and the *lga* BGC contigs. A dotted line is provided for easy comparison to find MAG(s) that have coverages within the range of the BGC coverage.

Reads were mapped back to all scaffolds and connections between contigs were inferred and counted using pair-end mapping. During the binning process, all contigs smaller than 3000 bp were discarded. Therefore any connected contigs with a length less than 3000 bp could not be used to determine if clustering of the BGC contigs into bin Loki_2 was correct. Of the contigs that carried fragments of the BGCs on them, only three were mapped to a total of six other contigs (Table M2). Of the six connected contigs, four belonged to Loki_2 and 2 belonged to Loki_1. The connections, taken in combination with the shared coverage of the Loki_2 bin allowed us to conclude that the BGC contig had been binned correctly. However, as the BGC contig was predicted to be connected to several contigs within this MAG, we cannot discount the possibility that the BGC has been shunted around following genomic rearrangement in closely related strains represented by this MAG. NODE_46_length_35697_cov_543.659124 was also noted as being potentially connected to the Loki_1 MAG. NODE_46 is present on one of the terminal edges of the “large section” of the *lga* BGC recovered from this metagenome. It is therefore possible that the connections are made with regions of the contig flanking the *lga* BGC.

**Table M2.** Number of predicted connections shared between the contig with the *lga* BGC and other binned contigs in the *L. okinawana* sample

| BGC contig | Connected contig | # connections | Connected contig/bin |
|---|---|---|---|
| NODE_180_length_12254_cov_531.844379 | NODE_46_length_35697_cov_543.659124 | 574 | BGC contig |
| NODE_161_length_14264_cov_611.590118 | NODE_347_length_6779_cov_822.969561 | 211 | Loki_2 |
| NODE_161_length_14264_cov_611.590118 | NODE_326_length_7089_cov_784.093697 | 54 | Loki_2 |
| NODE_161_length_14264_cov_611.590118 | NODE_264_length_8243_cov_871.029758 | 40 | Loki_2 |
| NODE_46_length_35697_cov_543.659124 | NODE_507_length_4709_cov_41.784758 | 20 | Loki_1 |
| NODE_161_length_14264_cov_611.590118 | NODE_446_length_5213_cov_891.608450 | 14 | Loki_2 |
| NODE_161_length_14264_cov_611.590118 | NODE_311_length_7428_cov_485.445246 | 8 | BGC contig |
| NODE_46_length_35697_cov_543.659124 | NODE_109_length_20869_cov_42.015487 | 7 | Loki_1 |

#### *L. rufipennis 2*

Binning via Autometa placed the contigs predicted to carry the lagriamide BGC, contigs NODE_27_length_17695_cov_1312.062096, NODE_62_length_13031_cov_1307.459395, NODE_241_length_7316_cov_1340.179445 and NODE_273_length_6877_cov_1327.437794 in bin Lruf2_2. Visualization of the dimension-reduced embedded 5-mer frequency data revealed that the contig was found at approximately the same abundance (coverage) as other contigs present in the Lruf2_2, and shared the same approximate 5-mer frequency pattern (Fig. M3)

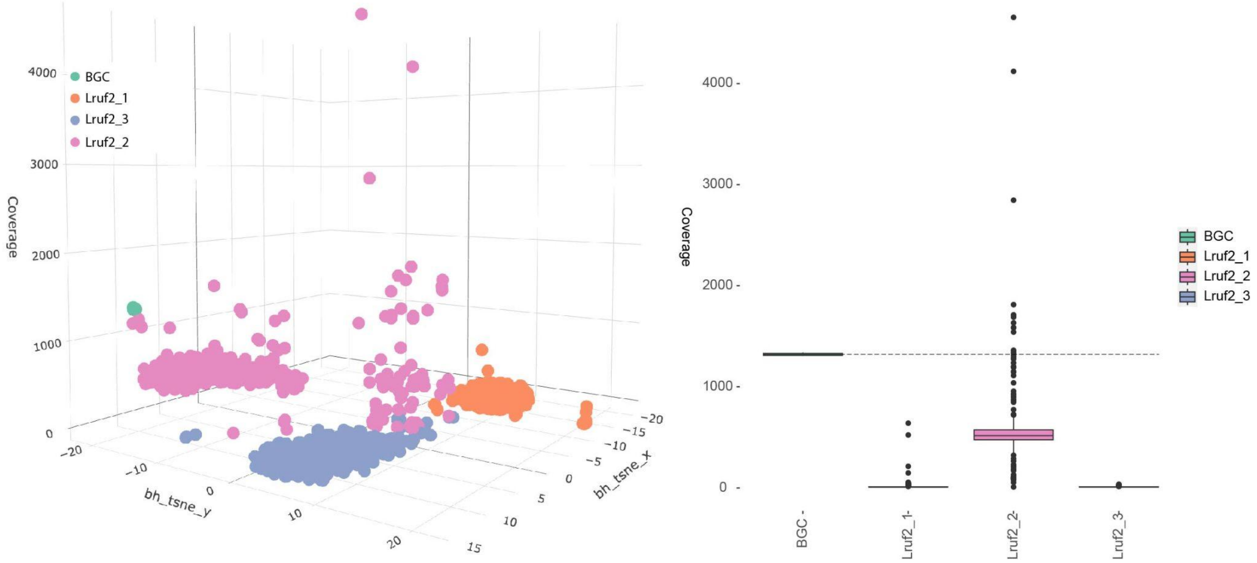

**Figure M3.** Placement of BGC contigs relative to MAGs. A) Three-dimensional visualization of all binned bacterial contigs from the *L. rufipennis 2* metagenome. Each point represents an assembled contig colored by the bin in which it was clustered. The contigs on which the lagriamide BGC was identified have been indicated. B) Range of contig coverages (a proxy for abundance) of each MAG and the *lga* BGC contig. A dotted line is provided for easy comparison to find MAG(s) that have coverages within the range of the BGC coverage.

Reads were mapped back to all scaffolds and connections between contigs were inferred and counted using pair-end mapping. During the binning process, all contigs smaller than 3000 bp were discarded. Therefore any connected contigs with a length less than 3000 bp could not be used to determine if clustering of the BGC contigs (NODE_27_length_17695_cov_1312.062096, NODE_62_length_13031_cov_1307.459395, NODE_241_length_7316_cov_1340.179445 and NODE_273_length_6877_cov_1327.437794) into Lruf2_2 was correct. Two of the contigs (NODE_27 and NODE_241) with fragments of the *lga* BGC were found to have possible pair-end-guided connections to a total of 4 contigs (Table M3). All four connected contigs were binned in Lruf2_2. These connections in combination with the matching coverage of the BGC carrying contigs and the bin into which they were clustered allowed us to be confident that the BGC had been binned correctly. We did note, however, that the average coverage of the BGC was approximately twice that of the average coverage of Lruf2_2, which could indicate that this genome has two copies of the BGC. This is further supported by the fact that there are at least four different contigs predicted to be connected to the BGC.

**Table M3.** Number of predicted connections shared between the contig with the *lga* BGC and other binned contigs in the *L. rufipennis 2* sample

| BGC contig | Connected contig | # connections | Connected contig bin |
|---|---|---|---|
| NODE_27_length_17695_cov_1312.062096 | NODE_1136_length_3410_cov_555.556556 | 303 | Lruf2_2 |
| NODE_27_length_17695_cov_1312.062096 | NODE_639_length_4632_cov_562.795170 | 83 | Lruf2_2 |
| NODE_27_length_17695_cov_1312.062096 | NODE_77_length_12150_cov_539.721859 | 28 | Lruf2_2 |
| NODE_241_length_7316_cov_1340.179445 | NODE_62_length_13031_cov_1307.459395 | 9 | Lruf2_2 |

#### *L. hirta* HG

Binning via Autometa placed the contig predicted to carry the lagriamide BGC, contig NODE_47_length_88863_cov_811.439765 from the hybrid assembly in bin LhHG_2. Visualization of the dimension-reduced embedded 5-mer frequency data revealed that the contig was found at approximately the same abundance (coverage) as other contigs present in the LhHG_2, and shared the same approximate 5-mer frequency pattern (Fig. M4)

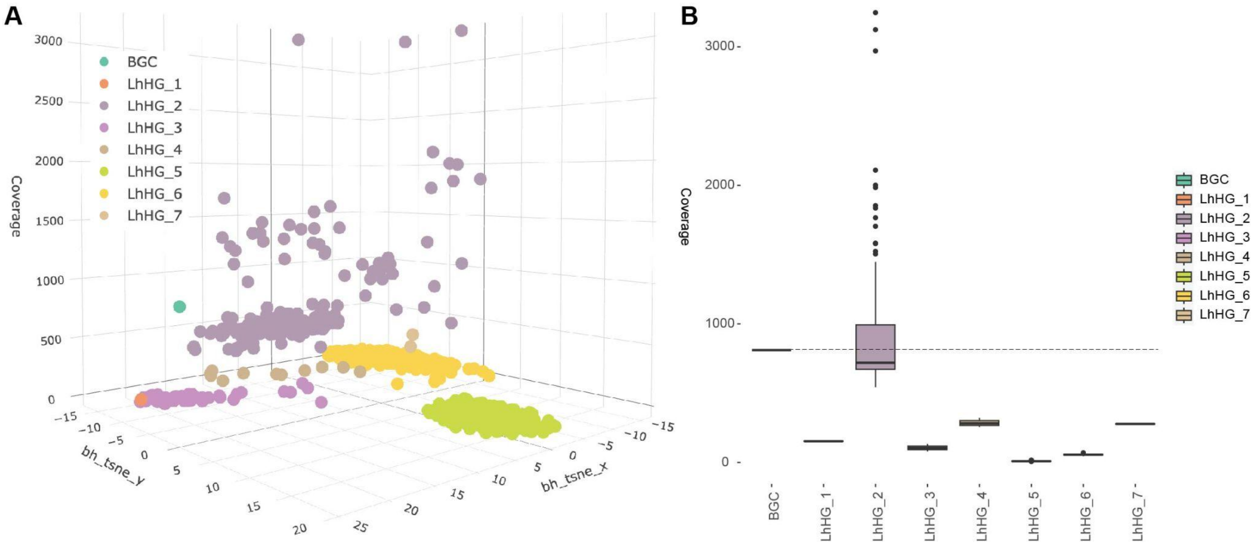

**Figure M4.** Placement of BGC contigs relative to MAGs. A) Three-dimensional visualization of all binned bacterial contigs from the *L. hirta* HG metagenome. Each point represents an assembled contig colored by the bin in which it was clustered. The contigs on which the lagriamide BGC was identified have been indicated. B) Range of contig coverages (a proxy for abundance) of each MAG and the *lga* BGC contig. A dotted line is provided for easy comparison to find MAG(s) that have coverages within the range of the BGC coverage.

Reads were mapped back to all scaffolds and connections between contigs were inferred and counted using pair-end mapping. During the binning process, all contigs smaller than 3000 bp were discarded. Therefore any connected contigs with a length less than 3000 bp could not be used to determine if the clustering of the BGC contig (NODE_47_length_88863_cov_811.439765) in bin LhHG_2 was correct. NODE_47 carried the *lga* BGC and was predicted to be connected with 113 different contigs, the ten contigs with the highest number of connections are provided in Table M4. Of the predicted connected contigs, 88 were binned in LhHG_2, 9 were binned in LhHG_3, 4 were binned in LhHG_4 and 12 were unclustered. As the majority of connected contigs were in LhHG_2, where the BGC contig had originally been binned, we were confident that the BGC had been assigned to the correct bin. However, once again, we cannot discount the possibility of genomic rearrangement variants due to several predicted connected contigs within the LhHG_2 MAG. Similarly to that observed in the L.okinawa sample, the contig from LhHG_4 predicted to be connected to the BGC may be due to a common sequence of the genes flanking the BGC that are found in non-lga-carrying variants.

**Table M4.** Top ten contigs connected to the lga BGC contig in the *L. hirta* HG sample by measure of the number of connections shared

| BGC contig | Connected contig | # connections | Connected contig bin |
|---|---|---|---|
| NODE_47_length_88863_cov_811.439765 | NODE_14076_length_4310_cov_714.523742 | 4412 | LhHG_2 |
| NODE_47_length_88863_cov_811.439765 | NODE_15881_length_3514_cov_659.124236 | 1870 | LhHG_2 |
| NODE_47_length_88863_cov_811.439765 | NODE_14748_length_4010_cov_2970.342232 | 1761 | LhHG_2 |
| NODE_47_length_88863_cov_811.439765 | NODE_22_length_133714_cov_111.253096 | 1454 | LhHG_3 |
| NODE_47_length_88863_cov_811.439765 | NODE_11104_length_5986_cov_779.033508 | 1076 | LhHG_2 |
| NODE_47_length_88863_cov_811.439765 | NODE_39_length_93533_cov_257.105836 | 812 | LhHG_4 |
| NODE_47_length_88863_cov_811.439765 | NODE_31_length_113099_cov_380.562138 | 767 | unclustered |
| NODE_47_length_88863_cov_811.439765 | NODE_9728_length_6936_cov_717.090392 | 542 | LhHG_2 |
| NODE_47_length_88863_cov_811.439765 | NODE_16884_length_3130_cov_764.072060 | 539 | LhHG_2 |
| NODE_47_length_88863_cov_811.439765 | NODE_14510_length_4117_cov_719.904703 | 484 | LhHG_2 |

#### *L. hirta* G

Binning via Autometa placed the contig predicted to carry the lagriamide BGC, contig NODE_6_length_91830_cov_314.588181 in bin LhG_1. Visualization of the dimension-reduced embedded 5-mer frequency data revealed that the contig was found at approximately the same abundance (coverage) as other contigs present in the LhG_1, and shared the same approximate 5-mer frequency pattern (Fig. M5)

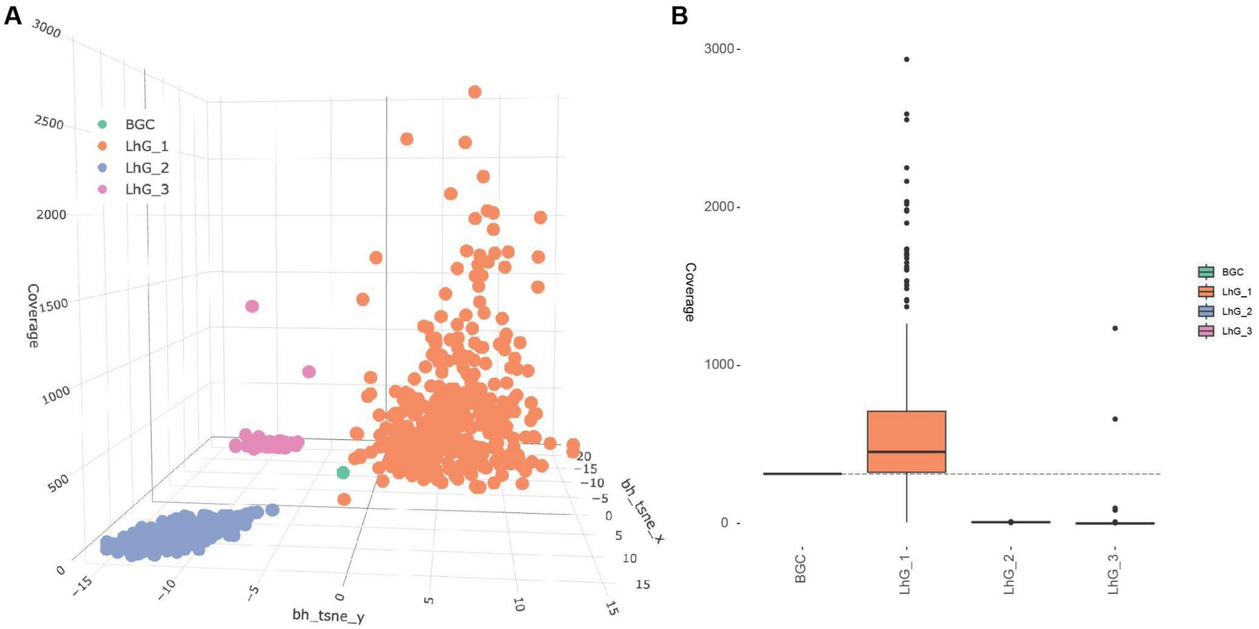

**Figure M5.** Placement of BGC contigs relative to MAGs. A) Three-dimensional visualization of all binned bacterial contigs from the *L. hirta* G metagenome. Each point represents an assembled contig colored by the bin in which it was clustered. The contigs on which the lagriamide BGC was identified have been indicated. B) Range of contig coverages (a proxy for abundance) of each MAG and the lga BGC contig. A dotted line is provided for easy comparison to find MAG(s) that have coverages within the range of the BGC coverage.

Reads were mapped back to all scaffolds and connections between contigs were inferred and counted using pair-end mapping. During the binning process, all contigs smaller than 3000 bp were discarded. Therefore any connected contigs with a length less than 3000 bp could not be used to determine if the clustering of the BGC contig (NODE_6_length_91830_cov_314.588181) in bin LhG_1 was correct. NODE_6 was predicted to be connected with 14 different contigs (Table M5). However, 13 of them were less than 3000 bp in length and therefore not binned. The only remaining connected contig shared only 20 connections had been binned in LhG_1, the same bin as NODE_6 which carried the BGC. While the connections are not conclusive, this in combination with the shared coverage of the LhG_1 bin allowed us to conclude that the BGC contig had been binned correctly.

**Table M5.** Number of predicted connections shared between the contig with the *lga* BGC and other contigs in the *L. hirta* G sample

| BGC contig | Connected contig | # connections | Connected contig bin |
|---|---|---|---|
| NODE_6_length_91830_cov_314.588181 | NODE_87_length_21318_cov_344.876551 | 20 | LhG_1 |
| NODE_6_length_91830_cov_314.588181 | NODE_1610_length_1447_cov_4694.536364 | 23 | Not binned, < 3000 bp |
| NODE_6_length_91830_cov_314.588181 | NODE_1028_length_1974_cov_232.890092 | 26 | Not binned, < 3000 bp |
| NODE_6_length_91830_cov_314.588181 | NODE_1891_length_1304_cov_1490.424809 | 34 | Not binned, < 3000 bp |
| NODE_6_length_91830_cov_314.588181 | NODE_67927_length_206_cov_1474.759494 | 56 | Not binned, < 3000 bp |
| NODE_6_length_91830_cov_314.588181 | NODE_67709_length_224_cov_25.061856 | 63 | Not binned, < 3000 bp |
| NODE_6_length_91830_cov_314.588181 | NODE_3058_length_984_cov_3299.152859 | 66 | Not binned, < 3000 bp |
| NODE_6_length_91830_cov_314.588181 | NODE_11568_length_533_cov_1099.406404 | 107 | Not binned, < 3000 bp |
| NODE_6_length_91830_cov_314.588181 | NODE_711_length_2655_cov_256.694620 | 140 | Not binned, < 3000 bp |
| NODE_6_length_91830_cov_314.588181 | NODE_67955_length_204_cov_4827.805195 | 161 | Not binned, < 3000 bp |
| NODE_6_length_91830_cov_314.588181 | NODE_68451_length_178_cov_4618.196078 | 188 | Not binned, < 3000 bp |
| NODE_6_length_91830_cov_314.588181 | NODE_67457_length_246_cov_61766.991597 | 211 | Not binned, < 3000 bp |
| NODE_6_length_91830_cov_314.588181 | NODE_1852_length_1320_cov_6490.212070 | 462 | Not binned, < 3000 bp |
| NODE_6_length_91830_cov_314.588181 | NODE_2247_length_1172_cov_31632.355024 | 851 | Not binned, < 3000 bp |

##### *L. hirta* SB

Binning via Autometa placed the contig predicted to carry the lagriamide BGC, contig NODE_38_length_73485_cov_188.551487 in bin LhSB_1. Visualization of the dimension-reduced embedded 5-mer frequency data revealed that the contig was found at approximately the same abundance (coverage) as other contigs present in the LhSB_1, but exhibited a divergent 5-mer frequency pattern from the other contigs in LhSB_1 (Fig. M6). This divergence is likely a result of the horizontal transfer of the BGC.

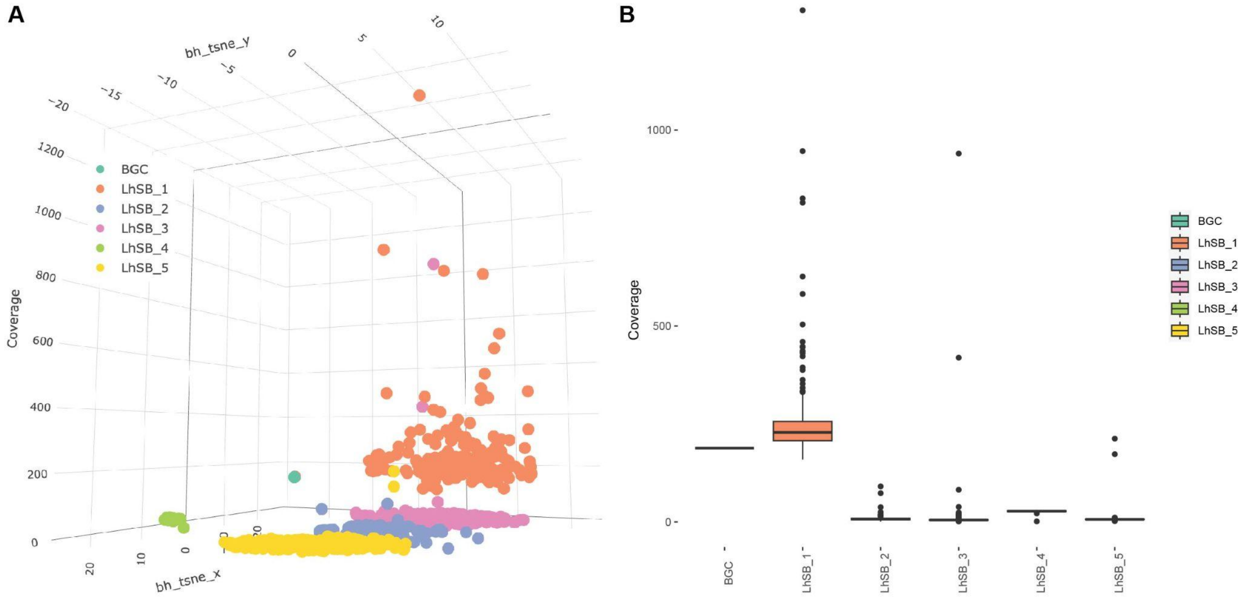

**Figure M6.** Placement of BGC contigs relative to MAGs. A) Three-dimensional visualization of all binned bacterial contigs from the *L. hirta* SB metagenome. Each point represents an assembled contig colored by the bin in which it was clustered. The contigs on which the lagriamide BGC was identified have been indicated. B) Range of contig coverages (a proxy for abundance) of each MAG and the *lga* BGC contig. A dotted line is provided for easy comparison to find MAG(s) that have coverages within the range of the BGC coverage.

Reads were mapped back to all scaffolds and connections between contigs were inferred and counted using pair-end mapping. During the binning process, all contigs smaller than 3000 bp were discarded. Therefore any connected contigs with a length less than 3000 bp could not be used to determine if the clustering of the BGC contig (NODE_38_length_73485_cov_188.551487) in LhSB_1 was correct. NODE_38 was connected to 17 contigs, 13 of which were less than 3000 bp and not binned, two of which were included in binning but unclustered, and two in LhSB_1 (Table M6). However, the two binned contigs shared the highest number of connections with NODE_38. While the connections are not conclusive, this in combination with the shared coverage of the LhSB_1 bin allowed us to conclude that the BGC contig had likely been binned correctly.

**Table M6.** Number of predicted connections shared between the contig with the *lga* BGC and other contigs in the *L. hirta* SB sample

| BGC contig | Connected contig | # connections | Connected contig bin |
|---|---|---|---|
| NODE_38_length_73485_cov_188.551487 | NODE_1533_length_3277_cov_446.117778 | 572 | LhSB_1 |
| NODE_38_length_73485_cov_188.551487 | NODE_416_length_11029_cov_1305.833609 | 370 | LhSB_1 |
| NODE_38_length_73485_cov_188.551487 | NODE_346425_length_248_cov_203.644628 | 173 | Not binned < 3000 bp |
| NODE_38_length_73485_cov_188.551487 | NODE_95658_length_651_cov_18721.959924 | 39 | Not binned < 3000 bp |
| NODE_38_length_73485_cov_188.551487 | NODE_120866_length_566_cov_331.615034 | 36 | Not binned < 3000 bp |
| NODE_38_length_73485_cov_188.551487 | NODE_268693_length_327_cov_219.820000 | 36 | Not binned < 3000 bp |
| NODE_38_length_73485_cov_188.551487 | NODE_712_length_5470_cov_2541.789257 | 28 | unclustered |
| NODE_38_length_73485_cov_188.551487 | NODE_1205_length_3738_cov_2389.862919 | 26 | unclustered |
| NODE_38_length_73485_cov_188.551487 | NODE_349186_length_230_cov_3007.213592 | 19 | Not binned < 3000 bp |
| NODE_38_length_73485_cov_188.551487 | NODE_2079_length_2850_cov_360.430040 | 19 | Not binned < 3000 bp |
| NODE_38_length_73485_cov_188.551487 | NODE_121539_length_564_cov_10854.48741 | 16 | Not binned < 3000 bp |
| NODE_38_length_73485_cov_188.551487 | NODE_359239_length_178_cov_433.901961 | 14 | Not binned < 3000 bp |
| NODE_38_length_73485_cov_188.551487 | NODE_345074_length_253_cov_158.714286 | 13 | Not binned < 3000 bp |
| NODE_38_length_73485_cov_188.551487 | NODE_276512_length_321_cov_0.592784 | 11 | Not binned < 3000 bp |
| NODE_38_length_73485_cov_188.551487 | NODE_91277_length_668_cov_395.173752 | 7 | Not binned < 3000 bp |
| NODE_38_length_73485_cov_188.551487 | NODE_71335_length_763_cov_9932.160377 | 6 | Not binned < 3000 bp |
| NODE_38_length_73485_cov_188.551487 | NODE_360825_length_164_cov_7200.216216 | 6 | Not binned < 3000 bp |

#### *L. grenieri*

Binning via Autometa placed the contig predicted to carry the lagriamide BGC, contig NODE_2_length_91929_cov_56.400046 in bin Lgren_6. Visualization of the dimension-reduced embedded 5mer frequency data revealed that the contig was found at approximately the same abundance (coverage) as other contigs present in the Lgren_6 bin, and shared the same approximate 5-mer frequency pattern (Fig. M7)

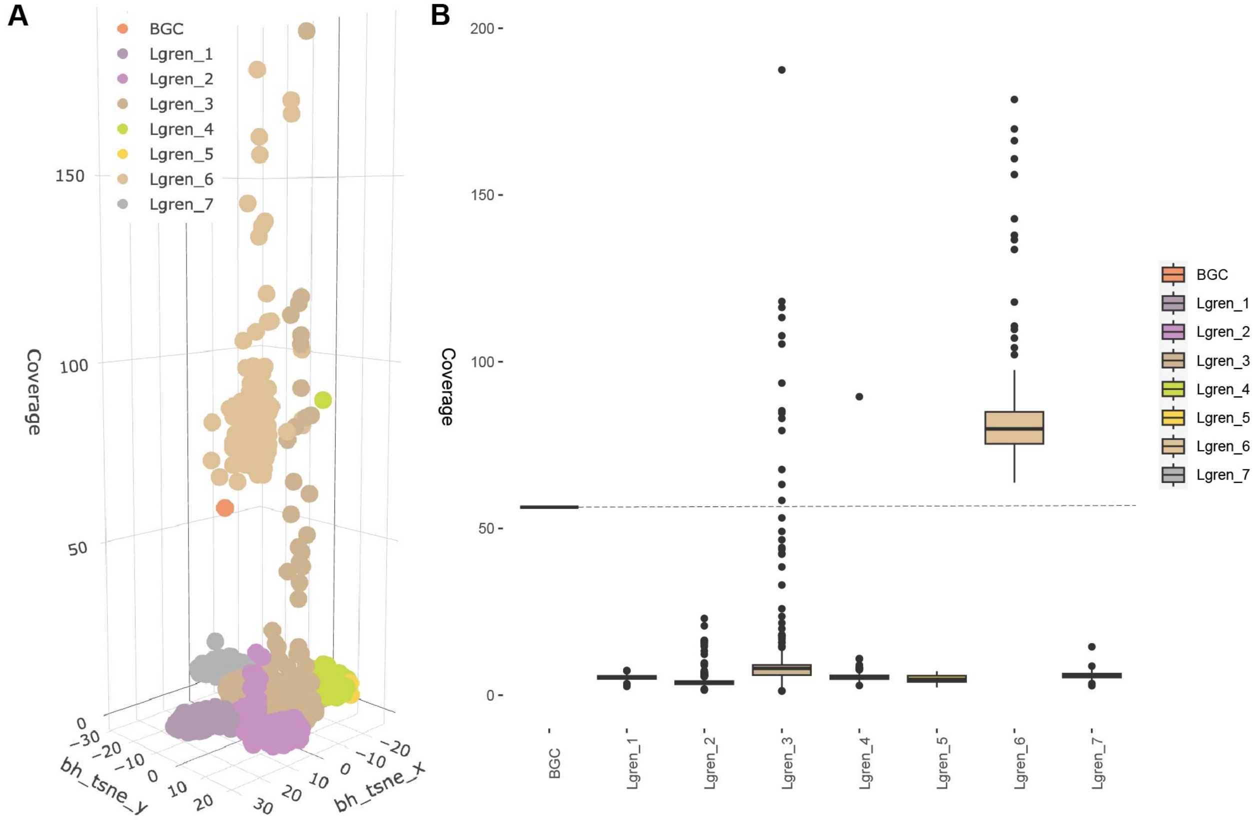

**Figure M7.** Placement of BGC contigs relative to MAGs. A) Three-dimensional visualization of all binned bacterial contigs from the *L. grenieri* metagenome. Each point represents an assembled contig colored by the bin in which it was clustered. The contigs on which the lagriamide BGC was identified have been indicated. B) Range of contig coverages (a proxy for abundance) of each MAG and the *lga* BGC contig. A dotted line is provided for easy comparison to find MAG(s) that have coverages within the range of the BGC coverage.

Reads were mapped back to all scaffolds and connections between contigs were inferred and counted using pair-end mapping. During the binning process, all contigs smaller than 3000 bp were discarded. Therefore any connected contigs with a length less than 3000 bp could not be used to determine if the clustering of the BGC contig (NODE_2_length_91929_cov_56.400046) in bin Lgren_6 was correct. NODE_2 was predicted to be connected with 21 contigs (Table M7) of which 18 were smaller than 3000 bp and not binned. The remaining three connected contigs had all been clustered into bin Lgren_6. We therefore concluded that the BGC contig, NODE_6, had been binned correctly.

**Table M7.** Number of predicted connections shared between the contig with the *lga* BGC and other contigs in the *L. grenieri* sample

| BGC contig | Connected contig | # connections | Connected contig bin |
|---|---|---|---|
| NODE_2_length_91929_cov_56.400046 | NODE_219914_length_451_cov_119.475309 | 56 | Not binned, < 3000 bp |
| NODE_2_length_91929_cov_56.400046 | NODE_219913_length_451_cov_153.567901 | 44 | Not binned, < 3000 bp |
| NODE_2_length_91929_cov_56.400046 | NODE_176878_length_554_cov_93.674473 | 26 | Not binned, < 3000 bp |
| NODE_2_length_91929_cov_56.400046 | NODE_384012_length_255_cov_343.367188 | 24 | Not binned, < 3000 bp |
| NODE_2_length_91929_cov_56.400046 | NODE_398287_length_224_cov_885.443299 | 19 | Not binned, < 3000 bp |
| NODE_2_length_91929_cov_56.400046 | NODE_7387_length_2898_cov_541.749910 | 15 | Not binned, < 3000 bp |
| NODE_2_length_91929_cov_56.400046 | NODE_156072_length_619_cov_108.682927 | 13 | Not binned, < 3000 bp |
| NODE_2_length_91929_cov_56.400046 | NODE_33637_length_1565_cov_98.745480 | 11 | Not binned, < 3000 bp |
| NODE_2_length_91929_cov_56.400046 | NODE_213063_length_465_cov_4910.715976 | 10 | Not binned, < 3000 bp |
| NODE_2_length_91929_cov_56.400046 | NODE_277500_length_360_cov_130.283262 | 9 | Not binned, < 3000 bp |
| NODE_2_length_91929_cov_56.400046 | NODE_159970_length_606_cov_59.947808 | 9 | Not binned, < 3000 bp |
| NODE_2_length_91929_cov_56.400046 | NODE_5592_length_3235_cov_71.177606 | 8 | Lgren_6 |
| NODE_2_length_91929_cov_56.400046 | NODE_84645_length_965_cov_83.837709 | 8 | Not binned, < 3000 bp |
| NODE_2_length_91929_cov_56.400046 | NODE_5333_length_3296_cov_1425.901862 | 7 | Not binned, < 3000 bp |
| NODE_2_length_91929_cov_56.400046 | NODE_48300_length_1319_cov_68.895973 | 7 | Not binned, < 3000 bp |
| NODE_2_length_91929_cov_56.400046 | NODE_2411_length_4634_cov_83.422010 | 7 | Lgren_6 |
| NODE_2_length_91929_cov_56.400046 | NODE_190627_length_517_cov_127.574359 | 7 | Not binned, < 3000 bp |
| NODE_2_length_91929_cov_56.400046 | NODE_226140_length_439_cov_79.423077 | 7 | Not binned, < 3000 bp |
| NODE_2_length_91929_cov_56.400046 | NODE_4365_length_3586_cov_110.713501 | 6 | Lgren_6 |
| NODE_2_length_91929_cov_56.400046 | NODE_384734_length_254_cov_368.645669 | 6 | Not binned, < 3000 bp |
| NODE_2_length_91929_cov_56.400046 | NODE_127531_length_727_cov_1732.205000 | 6 | Not binned, < 3000 bp |

#### *L. atripes*

Binning via Autometa placed the contigs predicted to carry the lagriamide BGC, contigs NODE_77_length_76027_cov_24.562306 and NODE_1122_length_8011_cov_22.845003 in bin

Latri_1. Visualization of the dimension-reduced embedded 5-mer frequency data revealed that the two contigs were found at the same abundance (coverage) as other contigs present in the Latri_1 bin, and shared the same approximate 5-mer frequency pattern (Fig. M8)

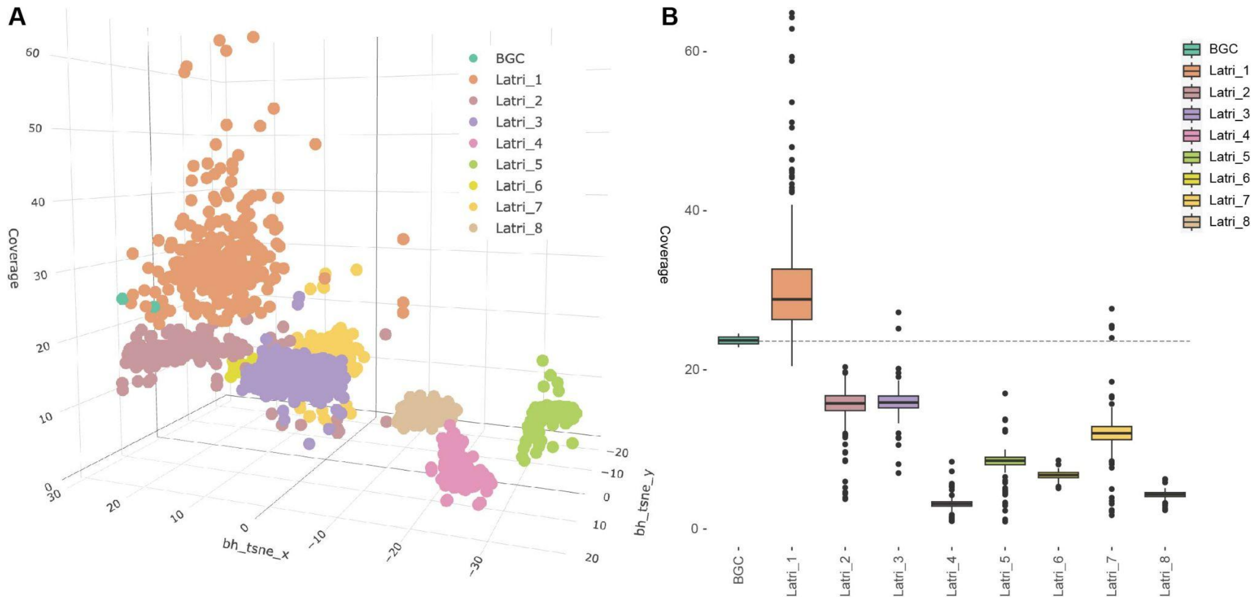

**Figure M8.** Placement of BGC contigs relative to MAGs. A) Three-dimensional visualization of all binned bacterial contigs from the *L. atripes* metagenome. Each point represents an assembled contig colored by the bin in which it was clustered. The contigs on which the lagriamide BGC was identified have been indicated. B) Range of contig coverages (a proxy for abundance) of each MAG and the *lga* BGC contigs. A dotted line is provided for easy comparison to find MAG(s) that have coverages within the range of the BGC coverage.

Reads were mapped back to all scaffolds and connections between contigs were inferred and counted using pair-end mapping. During the binning process, all contigs smaller than 3000 bp were discarded. Therefore any connected contigs with a length less than 3000 bp could not be used to determine if clustering of the BGC contigs (NODE_77_length_76027_cov_24.562306 and NODE_1122_length_8011_cov_22.845003) in bin Latri_1 was correct. In total, NODE_77 and NODE_1122 were predicted to be connected to only five other contigs (Table M8), three of which were too small to be binned. The binned contigs were assigned as unclustered and in Latri_1. While the connections are not conclusive, this in combination with the similar kmer-frequency (Fig. M8A) and similar average coverage of the Latri_1 bin (Fig. M8B) allowed us to conclude that the BGC contig had likely been binned correctly.

**Table M8.** Number of predicted connections shared between the contig with the *lga* BGC and other contigs in the *L. atripes* sample

| BGC contig | Connected contig | # connections | Connected contig bin |
|---|---|---|---|
| NODE_1122_length_8011_cov_22.845003 | NODE_2404_length_2697_cov_53.082101 | 89 | Not binned, < 3000 bp |
| NODE_77_length_76027_cov_24.562306 | NODE_2404_length_2697_cov_53.082101 | 64 | Not binned, < 3000 bp |

| | | | |
|---|---|---|---|
| NODE_77_length_76027_cov_24.562306 | NODE_1505_length_5161_cov_108.687525 | 23 | unclustered |
| NODE_77_length_76027_cov_24.562306 | NODE_860_length_11433_cov_58.832213 | 8 | Latri_1 |
| NODE_77_length_76027_cov_24.562306 | NODE_705465_length_243_cov_1342.2758 | 6 | Not binned, < 3000 bp |

#### *Ecnolagria* sp.

Binning via Autometa placed the contigs predicted to carry the lagriamide BGC, contigs NODE_1_length_71329_cov_79.783725, NODE_13080_length_6750_cov_78.631051 in bin Ecno_3. Contigs smaller than 3000 bp were not included in binning and therefore other contigs that carried fragments of the BGC were not visualized here. Visualization of the dimension-reduced embedded 5-mer frequency data revealed that the two contigs were found at the same abundance (coverage) as other contigs present in the Ecno_3 bin, and shared the same approximate 5-mer frequency pattern (Fig. M9)

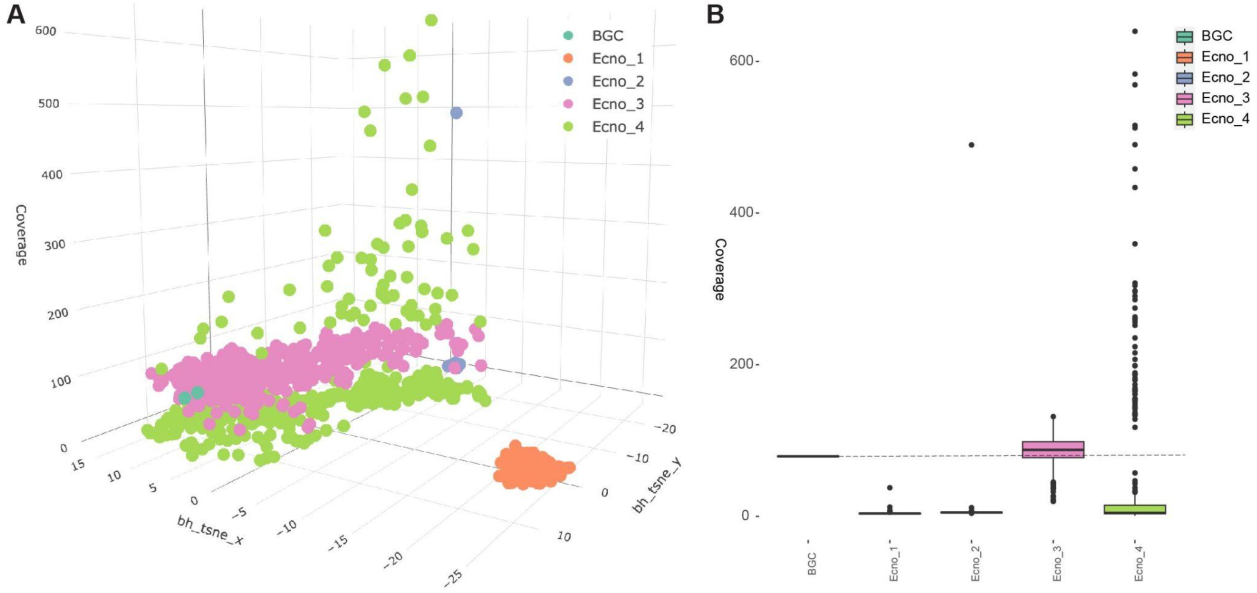

**Figure M9.** Placement of BGC contigs relative to MAGs. A) Three-dimensional visualization of all binned bacterial contigs from the *Ecnolagria* sp. metagenome. Each point represents an assembled contig colored by the bin in which it was clustered. The contigs on which the lagriamide BGC was identified have been indicated. B) Range of contig coverages (a proxy for abundance) of each MAG and the *lga* BGC contigs. A dotted line is provided for easy comparison to find MAG(s) that have coverages within the range of the BGC coverage.

Reads were mapped back to all scaffolds and connections between contigs were inferred and counted using pair-end mapping. During the binning process, all contigs smaller than 3000 bp were discarded. Therefore any connected contigs with a length less than 3000 bp could not be used to determine if clustering of the BGC contigs (NODE_1_length_71329_cov_79.783725, and NODE_13080_length_6750_cov_78.631051) in bin Ecno_3 was correct. Only a single connection was predicted using the pair-end mapping between NODE_1 and NODE_46491, which had been clustered into bin Ecno_3. While this alone was not substantial or convincing evidence that the BGC contig had been correctly assigned to Ecno_3, the corroborating coverage of the BGC contig and the Ecno_3 bin allowed us to conclude that the BGC likely had been binned correctly.

#### Prediction of domain functionality

The original *lga* BGC was identified in the genome of *Burkholderia* LvStB from the *L. villosa* 2019 sample [15]. Several domains were predicted to be non-functional due to mutations in key motifs. The antiSMASH algorithm similarly predicted several domains as non-functional. However, these were not in agreement with those predicted in the previous manuscript describing the *lga* BGC. Therefore, we extracted the amino acid sequences from all domains from all representative BGCs and aligned them using muscle (v5.1) [25].

The eleven representative BGCs spanned 13 different domain types: Acyl-carrier protein domains (ACP), Beta-branching acyl-carrier protein domains (ACP-beta), Adenylation domains (AMP-binding), Dehydratase domains (DH) and Dehydratase domain variant more commonly found in trans-AT PKS clusters (DHt), Condensation domains (C), Methyltransferase domains (cMT), Peptidyl-carrier protein domains (PCP), Enoylreductase domains (PKS_ER), Ketoreductase domains (PKS_KR), Ketosynthase domains (PKS_KS), Phosphopantetheine acyl carrier protein group (PKS_PP) and *Trans*-acyltransferase docking domains (*Trans*-AT_docking).

Full alignments proved too large to produce informative figures. All alignment files are available in Zenodo: 10.5281/zenodo.10381636.

##### Dehydratase domains

There were 90 dehydratase domains across the eleven *lga* BGCs. The four predicted DHt domains exhibited DAxxxxxxH motifs rather than the expected DSxxxxxW motif [26] which indicated that the differentiation into this sub-group was likely correct as none of the other DH domains included this motif. However, the conserved DH motifs HxxxGxxxxP, GYxYGPxF, DxxxQ/H, and LPFxW [27] are either extensively mutated or completely absent leading us to conclude that these domains may not be functional in all four DHt domains. Further, several mutations were present in the conserved motifs of the second DH domains in *lgaG* in all instances, in agreement with previous studies that predicted that this domain is likely non-functional. Unexpectedly, we discovered that the conserved HxxxGxxxxP motif is absent in the second DH of *lgaB* in the *L. villosa* 2023 sample, which may affect the functionality of the domain.

##### Ketosynthase domains

There were 193 ketosynthase domains across the eleven *lga* BGCs. The functionality of a KS domain can be predicted based on the catalytic triad CHH [28], present here in positions 187, 325, and 374 respectively. In the sixth KS domain of *lgaG*, in all *lga* BGCs recovered, the cysteine residue was mutated to a serine, the first histidine residue was mutated to an alanine, and the final histidine was intact. Similarly, the fifth KS domain of *lgaC* exhibited a mutation of the final histidine of the triad, which was instead mutated to a tyrosine in all cases. It was thus concluded that these two domains (*lgaG* KS6 and *lgaC* KS5) were likely non-functional.

#### Ketoreductase domains

There were 143 ketoreductase domains across the *lga* BGCs recovered from the eleven *Lagriinae* samples. The functionality of these domains is largely dependent on the conservation of the catalytic triad KSY [29]. The catalytic lysine (K) residue and the tyrosine (Y) residue were both present and correct in all sequences at position 119 and 162 respectively. The conserved serine residue was present at position 147 and appeared to have been mutated to an alanine in the third KR (KR3) in *lgaB* of all sequences, and it was concluded that these domains were likely non-functional.

#### Condensation domains

There were 22 condensation domains in the eleven recovered *lga* BGCs. The expected conserved motif of HHxxxDG [30], where only the second histidine and aspartate residues are required for catalytic function [30, 31] was present in all *lgaB* condensation domains but was HHxxxDA in the *lgaA* gene. This switch from glycine to alanine may have little effect due to their structural similarity and this motif variation has been observed in other functional condensation domains [32, 33], and as it has no known bearing on the catalytic ability of this domain, we presumed all 22 condensation domains to be functional.

#### Adenylation domains

There were 22 adenylation domains, approximately 394 residues in length, across the eleven recovered *lga* BGCs. Of the ten conserved motifs characteristic of adenylation domains [34], approximately 8 were found, with motifs A9 and A10 missing in all (Table M10). Critical motifs, A5 and A7 [35], in all *lga* adenylation domains are completely in keeping with expected motifs and we concluded that these domains were fully functional and predicted to bind to glycine.

**Table M10**. Expected conserved adenylation domain motifs relative to those identified in the *lga* BGC sequences. Matching residues are highlighted in green

| Motif | Identified | Position in alignment |
|---|---|---|
| A1: L(TS)YxEL | RWxYAEL | 20 - 26 |
| A2: LKAGxAYL(VL)P(LI)D | WQAGAAYVPLD | 66 - 76 |
| A3: LAYxxYTSG(ST)TGxPKG | LAYVLYTSGSTGRPKG | 145 - 160 |
| A4: FDxS | FDIA | 195 - 198 |
| A5: NxYGPTE | NLYGPTE | 285 - 291 |
| A6: GELxIxGxG(VL)ARGYL | GELYLGGAGLAQGGYH | 335 - 349 |
| A7: Y(RK)TGDL | Y(R/K)TGDL | 368 - 373 |
| A8: GRxDxQVKIRGxRIELGEIE | GRSDQQVKV | 386 - 394 |
| A9: LPxYM(IV)P | Not recovered | N/A |
| A10: NGK(VL)DR | Not recovered | N/A |

#### Enoyl reductase domains

There were 22 enoyl reductase domains across the eleven *lga* BGCs. We expected to find the glycine from the GxxxxxSxA motif, and the catalytic tyrosine and lysine residues from the Yxx(x)MxxxK motif intact [36] in all sequences to assume full functionality. We found the GxxxxxSxA motif as GDxVxx(I/L)A, and the Yxx(x)MxxxK motif as YIEIAMTALK, in all sequences.

As the required residues were present where expected we concluded that all enoyl reductase domains were likely functional.

#### Methyltransferase domains

Methyltransferase domains have at least 3 signature motifs [37] (Table M11), which were all identified in the methyltransferase domains in the *lga* BGC. We therefore concluded that all methyltransferase domains were likely functional.

**Table M11**. Expected conserved methylatransferase domain motifs relative to those identified in the *lga* BGC sequences

| Motif | Identified | Position in alignment |
|---|---|---|
| Motif 1: xL**E**(I/V)**G**x**G** | xLE(V/I)GAGTGGT | 40 - 51 |
| Motif 2: No single conserved residue | syDivvAAxxxLHAT | 112 - 118 |
| Motif 3: LLxxx**G**xxxx | LLxGWW | 163 - 168 |

#### Carrier protein domains

Carrier protein domains include peptide-carrier protein domains (PCP), acyl-carrier protein domains (ACP), beta-branching acyl-carrier protein domains (ACP-beta), and phosphopantetheine acyl carrier protein groups (PKS_PP). In total, there were 187 carrier protein domains present in the eleven *lga* BGCs. The majority of carrier proteins had a conserved serine residue present at position 85 of the alignment and due to its location, was assumed to be the serine required for phosphopantetheine attachment [38, 39]. The only two domains to not have this conserved serine were the two additional PKS_PP domains found in the *lgaC* gene from the *L. grenieri* and *L. okinawana* samples. Therefore these two were considered likely non-functional and all other carrier domains likely functional.
